# An egocentric map prioritizing peri-personal space in the mouse rostro-lateral visual area

**DOI:** 10.64898/2026.08.25.747123

**Authors:** Maria J. Sosa, Shayla Brooks, Maximillian Bluhm, Eliane Lei, Jean-Paul Noel

**Affiliations:** Department of Neuroscience, University of Minnesota, Minnesota, USA; Minnesota Robotics Institute, University of Minnesota, Minnesota, USA; Masonic Institute for the Developing Brain, University of Minnesota, Minnesota, USA

**Author notes:** Corresponding Author: Jean-Paul Noel. Equal contribution (Maria J. Sosa, Shayla Brooks).

## Abstract

All physical interactions between an organism and its environment occur within the space immediately adjacent to and surrounding its body, its peripersonal space (PPS). This space has been extensively studied behaviorally in humans, and through sparse single-neuron recordings in primates. However, how PPS is represented and organized at cellular and circuit scales remains poorly understood. Here, using dense extracellular recordings in the mouse rostro-lateral visual cortex (VISrl; >19,000 single units), we reveal the cellular and circuit organization of PPS in mice. Visuo-tactile neurons prioritize near-body space while also representing farther space in a direction-selective manner, tracking approaching but not receding objects across the environment. VISrl PPS neurons integrate vision and touch nonlinearly, and their tactile responses are progressively facilitated as visual objects near the body. PPS neurons are embedded in structured networks characterized by “like-to-like” functional connectivity and remap according to recent visuo-tactile statistics. Together, these findings establish VISrl as a circuit-accessible substrate for PPS, and reveal how near-body space is represented by a dynamic, plastic, multisensory cortical network.

## Introduction

A species’ evolutionary fitness depends on its ability to negotiate interactions with the external environment. These interactions, including appetitive or aversive physical contact and opportunities for action, necessarily unfold within the space immediately adjacent to and surrounding our bodies: our peri-personal space (PPS).

Paralleling the evolutionary importance of this space, seminal single-cell recordings in macaques demonstrated that the primate brain has a dedicated fronto-parietal network for encoding PPS (see^1–3^ for reviews). Indeed, neurons in the ventral intraparietal area (VIP), parietal area 7b, the ventral premotor cortex, and the putamen respond to touch on the body, as well as to visual stimuli presented in close spatial proximity to the body^4–8^. The visual receptive fields of these PPS neurons are anchored to specific body parts rather than the retina^9–11^, are depth-limited^12,13^, preferentially respond to approaching rather than receding stimuli^12^, and expand with increased speed of incoming visual stimuli^12,14^.

Building on these early neurophysiological observations, over the past three decades neuropsychological^15,16^, psychophysical^17^, electrophysiological^18–20^, and neuroimaging^21–24^ studies have detailed the dynamic and plastic properties of PPS in humans. For instance, PPS boundaries can expand after tool use^25,26^, shift with action demands or threats^27–30^, and reflect the recent statistical structure of our environments^31–33^. This work has also been accompanied by the development of biologically inspired neural networks^34–37^ and normative^38,39^ computational models (see^40^ for a review) which account for varied properties of PPS neurons and argue for distinct functional roles, from rapid and defensive motor responses^41^, to the prediction of future touch^39, 42, 43^, and computing the value of potential actions^44,45^.

Despite this progress, the cellular and circuit organization of PPS remains poorly understood. Classic neurophysiological studies^4–13^ were necessarily limited to relatively small numbers of neurons recorded one at a time, without access to population dynamics, putative cell types, or functional connectivity. As a result, we do not know how near-body space is represented across neural populations defined with millisecond and cellular resolution. We do not know whether PPS neurons are embedded in distinct excitatory and inhibitory networks, how PPS responses relate to representations of farther visual space, or how single neurons encoding near multisensory space adapt with changing demands or sensory statistics. In addition, while much of the human work delineating PPS fields relies on a nonlinear tactile facilitation during body-proximal presentations of auditory or visual cues^17,46–54^, this effect has not been described in PPS neurons. Indeed, although PPS neurons clearly converge visual and tactile information, they have not been shown to integrate visuo-tactile signals in the strict sense that the visuo-tactile response differs from the algebraic sum of visual and tactile responses. Work often cited as evidence that PPS neurons are multisensory^55^ demonstrated that neurons in VIP, a well-known site of PPS responses, integrate visuo-tactile signals. However, those neurons were not identified as PPS neurons.

Here, we sought to address the above-mentioned foundational gaps in our understanding of the neural circuitry underpinning PPS by establishing the mouse as a circuit-accessible model for studying PPS encoding. We focused on the rostro-lateral visual area (VISrl) because this area receives visual and tactile input^56^, is causally involved in visuo-tactile behavior (see^56,57^), and is selective among higher-order visual areas for binocular disparities corresponding to near space^58^. We hypothesized that VISrl may contain neurons with the canonical features of PPS cells, providing an entry point for studying how near-body space is represented, organized, and integrated across the senses, spatial reference frames, and scales.

We find that mouse VISrl harbors PPS neurons that respond to touch on the body and to nearby visual stimuli, prioritizing the space close to the body, but also coding for farther distances in a direction-selective manner: tracking objects that approach while discarding those that recede beyond reach. These neurons combine touch and vision non-linearly, their receptive fields expand with stimulus speed, their tuning adapts to the statistics of the environment, and they are embedded in a structured network demonstrating “like-to-like” functional connectivity. A subset of these neurons carry a grid-like code that hints at a bridge between neocortical and hippocampal representations of space.

## Results

### Peri-Personal Space Neurons in the Mouse Rostro-Lateral Visual Area

Mimicking established protocols to measure PPS fields in humans^17,46–54^, in a first experiment we presented head-fixed mice (n = 6 mice, n = 24 sessions) with an approaching or receding full-contrast white annulus (3 cm wide; presented on a 122 × 70 cm flat TV screen laying near parallel to the ground) traveling between 0 cm and 80 cm in depth, at either 50cm/s or 100cm/s (**Fig. 1A**). The visual stimuli onset and offset 1 second prior to or after moving, as to separate transient visual responses to on- and off-sets from putative visual depth receptive fields. In addition to these visual-only trials (n = 60 trials/session/visual condition), we also delivered tactile-only trials (n = 60 trials/session) via bilateral air-puff stimulation (50ms duration) on the animals’ vibrissa. Lastly, in visuo-tactile trials (n = 60 trials/session/condition) we combine the abovementioned visual stimulation with air-puffs delivered when the visual cue was at 1, 2, 4, 6, 8, 10, 12.5, 25, 50, or 75cm from the body. We simultaneously inserted two Neuropixel probes (NP1.0, 38 insertions) bilaterally to record from a total of 4966 single units in rostro-lateral visual area (VISrl; **Fig. 1B**; see **Fig. S1** and **Methods** for histology and probe localization). In addition, as off-targets we recorded neurons in CA1 (n = 2840), the dentate gyrus (n = 1684), primary visual cortex (n = 362), and the lateral posterior nucleus of the thalamus (n = 784), among other.

**Figure 1.**
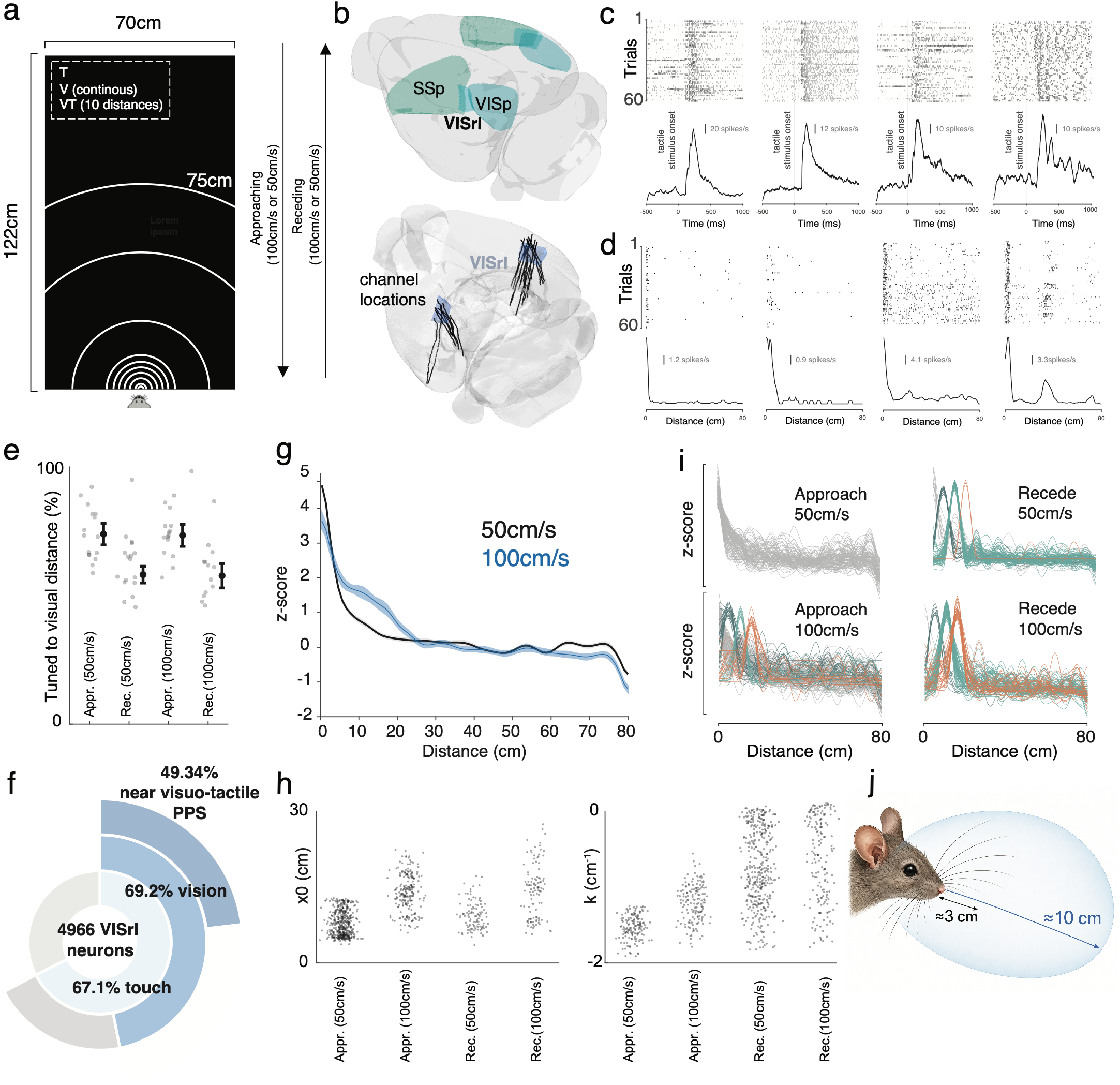
Peri-Personal Space Neurons in the Mouse Rostro-Lateral Visual Area. **A.** Schematic of the apparatus. **B.** Location of VISrl relative to primary visual (VISp) and somatosensory (SSp) cortices, and reconstruction of probe insertions (black) in Experiment 1 plotted on the common atlas of the Allen Institute. **C.** Example neurons responding to tactile stimulation. Top panel shows raster plots (each row is a trial, each dot is a spike) and bottom panel shows peri-stimulus time histograms (PSTHs). **D.** Example neurons tuned to visual distance. At difference from (**C**), the x-axis here is distance and not time. **E.** Fraction of tactile responsive neurons that are tuned to visual distance, as a function of direction (approach vs. recede) and speed (100cm/s vs. 50cm/s). Semi-transparent dots are individual sessions and opaque dots with error bars are mean and S.E.M. across sessions. **F.** Sunburst diagram showing how neurons were determined to be VISrl-PPS neurons; number of neurons in VISrl, fraction responding to touch, fraction modulated by visual distance, and finally fraction preferring near distances. **G.** Average visual tuning function for proximally-tuned visual neurons as a function of speed (50cm/s in black and 100cm/s in blue). Shaded areas are S.E.M. **H.** Left: Inflection point (x₀) for all VISrl PPS neurons as a function of visual direction and speed. Right: Slope (k) for these same neurons as a function of direction and speed. **I.** Normalized tuning functions for the top 300 neurons by mutual information tuned to visual distance, as a function of speed (top = 50cm/s; bottom = 100cm/s) and direction (left = approach; right = recede). Neurons with distance preferences not peaking at zero are colored. **J.** Schematic of mouse showing depth at which whiskers reach (∼3cm in depth) and the estimated size of their PPS fields (∼10cm, though as demonstrated, this PPS field is modulated by external factors such as the speed of approaching and receding stimuli).

Sixty-seven percent (mean ± S.E.M across animals; 67.1% ± 1.2%) of neurons in VISrl responded to air-puff stimulation (see **Fig. 1C** for examples; Wilcoxon signed-rank test, p < 0.05, mean firing rate -500ms to -50ms vs. 50ms to 500ms post-stimulus onset). This fraction was significantly higher than other brain regions recorded, except for well-known somatosensory areas such as the secondary whisker somatosensory thalamus, the anterior pretectal nucleus, and the superior colliculus (all other p < 0.05; **Fig. S2**). All subsequent analyses were restricted to VISrl neurons responsive to air-puff stimulation on the vibrissa. Of these tactile responsive units, 69.2% (S.E.M = 3.2%) were also visuo-spatially tuned (see **Fig. 1D** for examples; p < 0.05, permutation testing). Most of these units had a single peak in their distance tuning functions, but some had secondary fields (e.g., right-most neuron in **Fig. 1D**, showing a primary peak at a distance near 0cm from the body, but also secondary peaks at ∼35cm and ∼72cm). A greater fraction of neurons demonstrated tuning to distance when visual stimuli approached (72.1% ± 8.31%) than when it receded (54.4% ± 7.05%, p = 0.008) from the body (**Fig. 1E**). We did not observe a significant difference in the fraction of neurons tuned to visual distance as a function of speed (50cm/s vs. 100cm/s, p = 0.89). Forty-nine percent (49.34% ± 6.8%) of neurons responding to touch and being modulated by visual distance had visual tuning functions showing a preference for near distances (**Fig. 1F**, <20cm from the body; this criterion will be relaxed below). In other words, these are classically-defined PPS neurons: cells that respond to visual and tactile stimuli, with a preference for near visual objects (n = 1137 classically-defined PPS neurons).

To further determine whether VISrl PPS neurons exhibited a hallmark property of macaque PPS neurons, we examined the shape of PPS fields as a function of the speed of approaching and receding visual stimuli^12^. On average, increasing stimulus speed appeared to have two effects on VISrl PPS neurons: firing rates began to increase at farther distances from the body, and peak responses relative to baseline were reduced (**Fig. 1G**). To quantify this pattern, we fit firing rates as a function of distance with a sigmoid and estimated each neuron’s inflection point and slope, corresponding to parameters x₀ and k in **Equation 2** (see **Fig. S3** for example fits). The distance bifurcating the near and far space was on average 8.02 cm from the body when visual stimuli approached at 50 cm/s, and expanded to 13.08 cm when stimuli approached at 100 cm/s (S.E.M. = 0.29 cm and 0.35 cm, respectively; p < 0.0001; **Fig. 1H**, left). This expansion parallels the behavior of macaque PPS neurons, whose receptive fields extend farther in depth as the speed of an approaching stimulus increases^12^.

Increasing stimulus speed also altered the shape of the distance tuning functions (**Fig. 1G**). The slope with which firing rate decayed as visuotactile distance increased was shallower in the 100 cm/s condition (-1.2 ± 0.36) than in the 50 cm/s condition (-1.58 ± 0.12; p = 0.033). More generally, slopes were shallower for receding stimuli than for approaching stimuli (p = 0.048; **Fig. 1H**, right). Together, these results suggest that faster visual motion expands and broadens the spatial extent of VISrl PPS responses. However, a striking feature of these data was the increased heterogeneity induced by faster stimulus speeds. To examine this variability, we focused on the 300 neurons with the strongest distance tuning, defined by mutual information (**Equation 1**), and visualized their tuning functions across speed and direction conditions. When stimuli approached at 50 cm/s, these neurons showed tuning functions that all peaked approximately 0 cm from the body (**Fig. 1I**, top left). However, when the speed increased to 100 cm/s, many of the same neurons exhibited peak responses at distances farther from the body (**Fig. 1I**, bottom left, colored; χ² test, p < 0.001). Thus, faster speeds did not simply expand PPS fields and weaken distance tuning. Instead, neurons shifted the preferred distance away from the body. A similar pattern was observed for receding stimuli. Even at 50 cm/s, a subset of neurons preferred distances slightly farther from the body, approximately 5 to 15 cm. At 100 cm/s, this pattern became more pronounced, with a larger fraction of neurons exhibiting preferred distances away from the body (χ² test, p = 0.003). Thus, although population averages demonstrate that increased speed produced a larger and less sharply defined PPS representation, single-neuron analyses revealed a more structured effect: many neurons that were most strongly tuned to immediate contact at lower speeds shifted their preferred distance outward at higher speeds.

Together, these findings show the existence of PPS neurons in mouse VISrl: neurons that respond to tactile and visual stimuli, are modulated by visual proximity to the body, preferentially respond to approaching rather than receding stimuli, and exhibit visuotactile receptive fields whose depth is modulated by the speed of incoming visual cues. On average, these proximally tuned VISrl neurons had PPS fields that extended two to three times beyond the reach of the animals’ whiskers (respectively, ∼ 10 cm vs. ∼3cm; **Fig. 1J**).

### Prioritizing the Near Space, but also Coding Far Space in a Direction-Selective Manner

The analyses above demonstrate the existence of classically defined PPS neurons in mouse VISrl. However, this analysis relied on predetermined criteria to identify proximity-tuned visual and tactile neurons, arbitrarily operationalized as visual responses peaking within 20 cm of the body. Approximately half of the visual and tactile-responsive neurons in VISrl met these criteria, leaving the remaining neurons unexplained.

To account for the full population of visuo-tactile neurons, we took a data-driven approach by z-scoring all distance-dependent response profiles and clustering them using k-means, selecting the number of clusters that maximized the mean silhouette score. This metric quantifies how well each neuron fits within its assigned cluster relative to other clusters. For visualization, we then projected all normalized visual tuning functions onto a two-dimensional, nonlinear latent space using t-SNE^59^ (**Fig. 2A**; example shown for approaching stimuli at 100 cm/s). This procedure again suggested that proximity tuning may be best understood as a series of discrete basis functions rather than as a continuum. Across the four visual motion conditions tested, approaching at 50 or 100 cm/s and receding at 50 or 100 cm/s, neurons clustered into 4 to 6 groups. **Figure 2B** shows the mean tuning function and example neurons for each cluster when visual cues approached or receded from the mouse body at 100 cm/s. In the case of approaching visual cues, the most proximal neurons peaked at a distance of 0 cm from the body (**Fig. 2B**, orange, top). These were the most numerous neurons, representing 38.2% of all visuo-tactile neurons in VISrl during 100 cm/s approach. Other clusters had visual tuning functions that peaked at 26.0 cm (blue), 43.4 cm (green), 53.9 cm (yellow), and 62.6 cm (purple) from the body.

**Figure 2.**
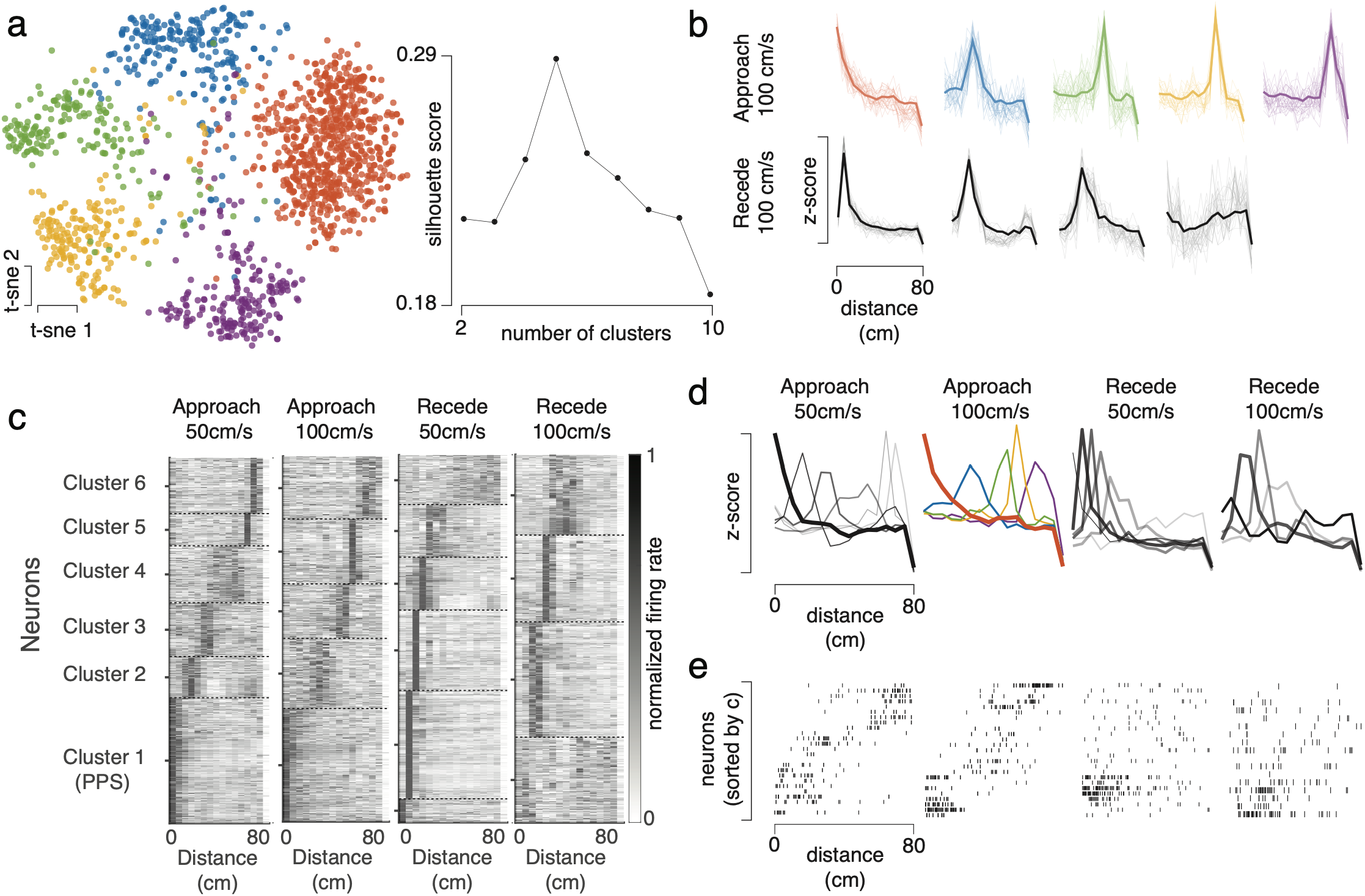
Prioritizing the Near Space, but also Coding Far Space in a Direction-Selective Manner. **A.** Left: Two-dimensional projection (t-sne) of normalized visual tuning functions, color-coded by cluster assignment. Right: Silhouette score as a function of number of clusters, peaking at 5 clusters. This data is for the approach at 100cm/s data. **B.** Mean (opaque and thick) and example (semi-transparent) visual tuning function for each of the clusters identified for the approach and recede at 100cm/s conditions. **C.** Heatmaps showing the normalized firing rate for each neuron (y-axis) and visual motion condition. Neurons are sorted by cluster assignment, with increasing cluster number indicating preferences for farther distances. Dashed lines separate clusters. **D.** Mean tuning function for each cluster (overlayed), and visual motion condition. Importantly, the frequency of each cluster is indicated by the thickness of the curves. Tuning functions with very proximal preferences are most common, and far spaces are only represented when visual cues approach the body, but not when they recede from it. **E.** Four example single trials (1 per visual motion condition, in the same order as in **D**). Neurons are sorted by cluster assignment, but not by their individualized preferred visual distance.

A similar pattern was observed across the other visual motion conditions (**Fig. 2C**). Neurons segregated into clusters that were well defined by their preferred visuo-tactile distance. For approaching visual cues, preferred distances tiled the full range of distances tested, with the most common preference being nearest to the body (as in **Fig. 2B**; χ² test, p < 0.0001). The relative fractions of the other preferred distances were approximately uniform (χ² test omitting Cluster 1, p = 0.19). In contrast, for receding visual cues, the most common preferred distance was one step away from the most proximal distance (10.4 cm for 50 cm/s and 16.7 cm for 100 cm/s). Even more strikingly, when visual cues receded from the body, there were no visuo-tactile neurons with preferences for the farthest distances tested. To illustrate this point, we overlaid the mean visual tuning functions for each cluster, motion direction, and velocity (**Fig. 2D**). The thickness of each curve represents its relative frequency. When stimuli approached the body, VISrl neurons appeared to hand off the stimulus representation from one distance-tuned population to the next, thereby tiling the full tested space. However, when visual stimuli receded from the body, VISrl neurons appeared to track stimulus position only through approximately the nearest half of the tested distances, up to about 40 cm. This effect was also visible on single trials when leveraging the full population of concurrently recorded visuo-tactile neurons (**Fig. 2E**, panel order is as in **Fig. 2D**: approach 50cm/s, approach 100cm/s, recede 50cm/s, and recede 100cm/s).

Together, these findings demonstrate that although VISrl prioritizes the encoding of PPS, it also represents farther distances in a direction-dependent manner. Specifically, VISrl appears to track the full trajectory of approaching stimuli, but only the behaviorally relevant near space when objects recede from the body. This effect was visible even on single trials when leveraging full simultaneously recorded populations. We return to population dynamics in the section after next.

### Virtual Touch and Modulation of Tactile Responses by Visual Distance

So far, we have examined touch-only and vision-only trials. These trial types are sufficient to identify PPS neurons as classically defined in the seminal macaque neurophysiology literature^4–13^. However, here we also included a third trial type: visuo-tactile trials in which touch was delivered when the visual cue was at different distances from the body. This design parallels the most established approach for indexing PPS in humans^17,46–54^, in which tactile facilitation, most commonly measured through reaction times, is quantified as a function of visual proximity. To our knowledge, this effect has not been demonstrated at the level of single neurons.

We first visualized single neurons by plotting raster plots (first ten columns) and peri-stimulus time histograms (PSTHs; right-most column) aligned to tactile onset, organized as a function of visuo-tactile distance (**Fig. 3A**). This analysis revealed neurons whose response amplitudes appeared to be modulated by visuo-tactile distance. More strikingly, several neurons exhibited two distinct evoked responses (**Fig. 3A**, rows 1 to 6). The first response is aligned to tactile onset, by design. The timing of the second response varied systematically with both visuo-tactile distance and visual stimulus velocity. Specifically, this second peak occurred when the approaching visual object reached the body, corresponding to a “virtual touch” event. Indeed, the time interval between the tactile-evoked response and the second response was longer when the visual stimulus was farther from the body at tactile onset, because a cue positioned at 75 cm requires more time to reach the body than a cue positioned at 10 cm, for a fixed velocity. Similarly, the interval between the two evoked responses for a given distance was longer for slower visual stimuli (**Fig. 3A**, top three rows, 50 cm/s) than for faster stimuli (**Fig. 3A**, next three rows, 100 cm/s). Lastly, this effect was absent for receding cues (**Fig. 3A**, rows 7 and 8), consistent with the absence of a future body-contact event, as indicated by the lack of vertical bars marking the timing of virtual touch. The presence of these secondary peaks is consistent with neurons that respond both to real touch on the body and to visual stimuli located very near the body. Thus, these neurons meet the criteria for PPS neurons while also suggesting their functional role in signaling not only physical contact, but also imminent or virtual contact.

**Figure 3.**
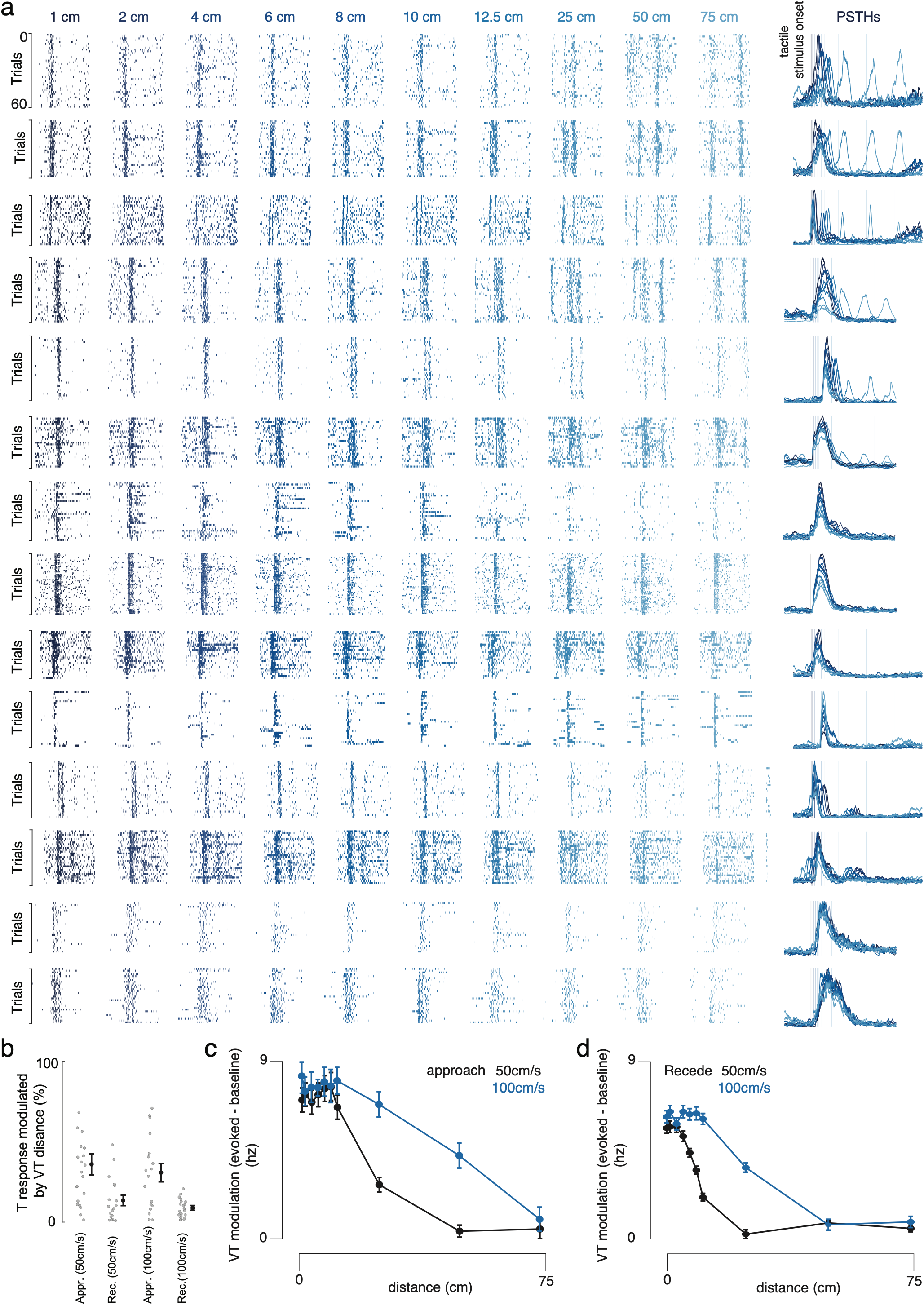
Modulation of Tactile Responses by Visual Distance. **A.** Raster plots and PSTHs for example neurons, as a function of visuo-tactile distance (darker blue = closer distance). Vertical bars in the right-most column (PSTHs) indicate when touch occurs and/or is implied to occur **B.** Fraction of neurons whose tactile response is modulated by visuo-tactile distance. Individual semi-transparent dots are means for a given session, and opaque black are means and S.E.M. across sessions. **C.** Evoked visuo-tactile response relative to the evoked tactile response (y-axis), as a function of visuo-tactile distance (x-axis) and visual velocity (black = 50cm/s, blue = 100cm/s). Error bars are S.E.M. **D.** As **C** for receding visual cues.

Not all neurons whose tactile responses were seemingly modulated by visual distance exhibited explicit secondary peaks corresponding to virtual touch (**Fig. 3A**, row 9). In addition, although most neurons showed enhanced tactile responses as visual cues approached the body (**Fig. 3A**, rows 7 to 9), some showed the opposite pattern, with tactile responses suppressed by visual proximity (**Fig. 3A**, row 10). Finally, in trials in which visual objects receded from the body, a small minority of neurons exhibited weak secondary peaks (**Fig. 3A**, rows 11 and 12). These responses did not depend on absolute visuo-tactile distance or velocity, but instead appeared to reflect the time elapsed since touch. Other visuo-tactile neurons showed no clear tactile facilitation as a function of visual proximity (**Fig. 3A**, rows 13 and 14). Overall, 30.2% (± 4.5%) of tactile-responsive neurons in VISrl showed tactile facilitation by visual proximity when visual cues approached the body, whereas 11.6% (± 3.6%) showed such facilitation during receding cues (p = 0.0008). The fraction of neurons whose tactile responses were modulated by visual distance was not strongly influenced by visual cue velocity (p = 0.63; **Fig. 3B**).

Next, we pooled all neurons showing significant tactile modulation by visuo-tactile distance. For each neuron, we computed the difference between the evoked response in visuo-tactile trials and the response in tactile-only trials. Evoked responses were defined as spike counts in the response window, 50 to 500 ms after tactile onset, minus spike counts in the baseline window, 500 to 50 ms before tactile onset. We then averaged these response differences across neurons. This analysis showed that, when visual cues approached the mouse at 50 cm/s, visuo-tactile responses did not differ from tactile-only responses at 50 or 75 cm from the body, but were significantly enhanced at distances below 25 cm (one-way ANOVA, p < 0.001; post hoc paired t-tests against zero, p < 0.01; **Fig. 3C**). When visual cues approached at 100 cm/s, visuo-tactile responses did not differ from tactile-only responses at 75 cm, but were significantly enhanced at all other distances tested (one-way ANOVA, p < 0.001). Interestingly, for both velocities, facilitation appeared to saturate once visual cues were within 12.5 cm of the body, such that responses at 1 cm did not differ from responses at 12.5 cm (all p > 0.56). A similar pattern was observed when visual cues receded from the body (**Fig. 3D**). At 50 cm/s, visuo-tactile responses were facilitated from 1 to 12.5 cm (all p < 0.001), whereas at 100 cm/s, facilitation extended up to 25 cm (all p < 0.001).

Together, these findings show at a cellular and millisecond resolution that tactile responses in VISrl are facilitated as a function of visuo-tactile distance, and that this facilitation depends on both motion direction and velocity. Moreover, the visualization of raster plots and PSTHs during approaching visual motion highlights a potential functional role for near-body visual receptive fields: VISrl PPS neurons respond not only to physical touch, but also to “virtual touch,” when approaching visual objects “make contact” with the body.

### A Population of Peri-Personal Space Neurons

On average, we simultaneously recorded from 22.17 VISrl-PPS neurons per session (range: 2 to 96). This enabled the first characterization of a network of PPS neurons, including the functional connectivity between excitatory and inhibitory cells, the probability of putative monosynaptic connections as a function of physical and feature distance between neurons (i.e., whether PPS neurons are organized according to “like-to-like” or “all-to-all” connectivity motifs), and their population dynamics.

We first examined the distribution of trough widths in extracellular spike waveforms (**Fig. 4A**, example session). Waveform widths were bimodally distributed, with most neurons exhibiting either narrow waveforms, with trough widths <0.35 ms, or broad waveforms, with trough widths >0.45 ms. We classified narrow-waveform units as putative inhibitory neurons (**Fig. 4A**, red) and broad-waveform units as putative excitatory neurons (**Fig. 4A**; see^60^ for a similar approach). Across sessions, 78.5% (SEM = 4.86%) of VISrl-PPS neurons were classified as putative excitatory, and 13.6% (SEM = 3.60%) were classified as putative inhibitory. Units with intermediate waveform widths were left unclassified.

**Figure 4.**
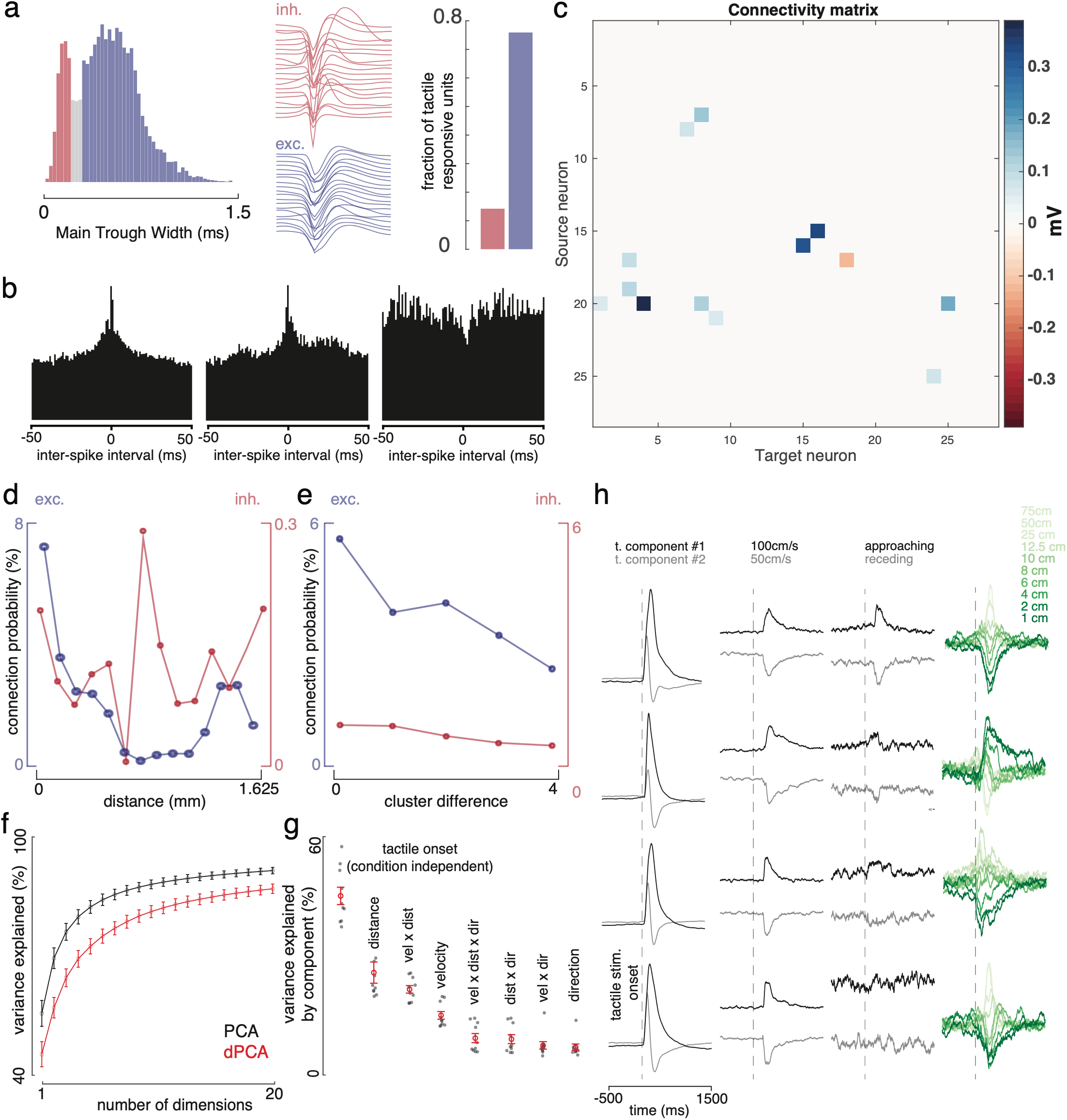
A Population of Peri-Personal Space Neurons. **A.** Example histogram of trough width of spike waveforms, clustered into narrow (red, putatively inhibitory neurons) and broad (blue, putatively excitatory neurons) waveforms, and their relative fraction (for example session). **B.** Three example inter-spike cross-correlograms (x-axis is time from -50 to 50ms and y-axis are counts). The first two depict likely excitatory connections (positive peak near zero ms) and the third depicts a likely inhibitory connection. **C.** Connectivity matrix from source neurons (y-axis) to target neurons (x-axis), with the color indicating connection strength (in mV, output from GLMCC). **D.** Connection probability as a function of distance between neurons and whether the connection is excitatory (blue, left y-axis) or inhibitory (red, right y-axis). **E.** Connection probability as a function of feature distance (i.e., difference in cluster number, when these clusters are sorted by peak location) and whether the connection is excitatory (blue, left y-axis) or inhibitory (red, right y-axis). **F.** Variance explained as a function of number of dimensions, for PCA (black) and dPCA (red). Error bars are S.E.M. **G.** Variance explained by the different subspaces. Black semi-transparent dots are individual sessions (all in VISrl) and red opaque dots and error bars are means and S.E.M across sessions. **H.** Neural data from four different sessions aligned to tactile stimulation (-500ms to 1500ms) and projected onto different subspaces: condition independent (touch), velocity, direction, and visuo-tactile distance. Right-most shows the visuo-tactile distance subspace, with gradients on green showing the distances tested.

Next, we applied a generalized linear model for cross-correlation analysis (GLMCC^61^) to infer putative monosynaptic interactions between simultaneously recorded VISrl-PPS neurons. For each directed neuronal pair, spike timing relationships were summarized as a cross-correlation function between −50 to 50 ms lags, at 1 ms resolution (**Fig. 4B**). A generalized linear model was then fit to isolate short-latency monosynaptic interactions from slower co-fluctuations in firing rate. Putative connections were classified as excitatory or inhibitory based on the sign of the inferred coupling term. These putative couplings were retained only when their estimated interaction exceeded the confidence interval for the null hypothesis of no connection and when a pre-trained convolutional neural network identified the pair as a putative monosynaptic connection^61,62^ (**Fig. 4C**, example session connectivity matrix). Overall, 74.3% of putative monosynaptic connections were classified as excitatory, and there was 90.1% agreement between waveform-based and spike-timing-based classifications of neuronal type.

Interestingly, the distance between neurons, both in physical space and in “feature space,” was a clear determinant of putative connectivity. For excitatory cells, up to 7.85% (SEM = 1.3%) of neurons were estimated to be coupled to other neurons within 116 μm (**Fig. 4D**). This probability initially decreased with distance, but then rebounded between 1392 and 1508 μm. Although inhibitory connections were less common overall (right y-axis in **Fig. 4D**), their probability peaked near the trough of the excitatory connection probability, about 800 μm from the reference neuron (**Fig. 4D**, red). This pattern is reminiscent of a Difference-of-Gaussians profile, with excitation, then inhibition, then excitation again, which has often been used to approximate lateral connectivity in biologically plausible neural network models of PPS^34–37^. Here, we provide an empirical estimate of this computational parameter.

We also computed the likelihood of putative coupling as a function of functional similarity (**Fig. 4E**). Specifically, we calculated connection probability as a function of the difference in cluster number. For example, two units assigned to cluster 1 in **Fig. 2C** would have a cluster difference of 0, whereas one unit assigned to cluster 1 and another assigned to cluster 4 would have a cluster difference of 3. Clusters were sorted according to their preferred visual distance. This analysis showed that excitatory units were most likely to couple with units encoding similar features, consistent with “like-to-like” connectivity (one-way ANOVA, p = 0.0012). In contrast, inhibitory units showed a relatively flat connectivity profile (p = 0.40). Together with the distance-dependent coupling profile shown in **Fig. 4D**, this result suggests the possible existence of a topographic map of visuo-tactile preferences in VISrl. A broad and undefined inhibitory tone coupled with “like-to-like” connectivity in excitatory connections is also suggestive of attractor dynamics.

Biologically, these unit-to-unit connectivity matrices should constrain the structure of population dynamics. Therefore, we next examined population-level state-space trajectories using demixed principal component analysis^63^ (dPCA). This method is conceptually similar to PCA, but preserves interpretability of the learned components, or subspaces, with respect to task features. In the current dataset, the first 20 PCs explained 94.5 % of the total variance, and dPCA explained 90.4% of the PCA-explained variance across the same number of components (**Fig. 4F**). The task-dependent feature explaining the most variance was visuo-tactile distance (25.6 %; see **Fig. S4** for other areas), followed by the interaction between velocity and distance (21.5%), velocity (15.1%), and the interaction among distance, velocity, and direction (9.62%; **Fig. 4G**). In total, visuo-tactile distance, including its main effect and all interactions in which it participated, accounted for 65.5% of neural variance in VISrl. **Figure 4H** depicts example subspaces encoding tactile stimulation, visual velocity, visual direction, and visuo-tactile distance. Each subspace is aligned to tactile stimulus onset. Notably, the velocity and direction subspaces were segregated before tactile onset, whereas the visuo-tactile distance subspace bifurcated after tactile onset and tiled a gradient from the nearest to the farthest tested distances (**Fig. 4H**, rightmost panel, green gradient).

### Peri-Personal Space Neurons Integrate Visuo-Tactile Signals

Next, we aimed to characterize the multisensory integration properties of PPS neurons. To this end, a new cohort of animals (n = 7 mice, n = 21 sessions) was first presented with an abbreviated version of the PPS mapping protocol from Experiment 1 (see above), which only included approaching stimuli at a single velocity of 125 cm/s. Animals were then presented with a separate routine designed to characterize multisensory integration more directly. In this protocol, visual stimuli did not approach or recede from the animal. Instead, on each trial, a visual annulus appeared for 50 ms at a fixed distance from the mouse, either alone or concurrently with a 50 ms tactile stimulus. Visual stimuli were presented at 1, 2, 4, 6, 8, 10, 12.5, 25, 50, or 75 cm from the mouse, and at 100%, 25%, or 12.5% contrast (**Fig. 5A**, a single visual distance shown). The protocol also included tactile-only trials. Because both protocols were performed within the same recording session, we first identify PPS neurons and then characterize the multisensory response properties of these same units. We recorded extracellular single-unit activity from VISrl across 38 probe insertions (3171 single units; see **Fig. S5A** for histological reconstructions).

**Figure 5.**
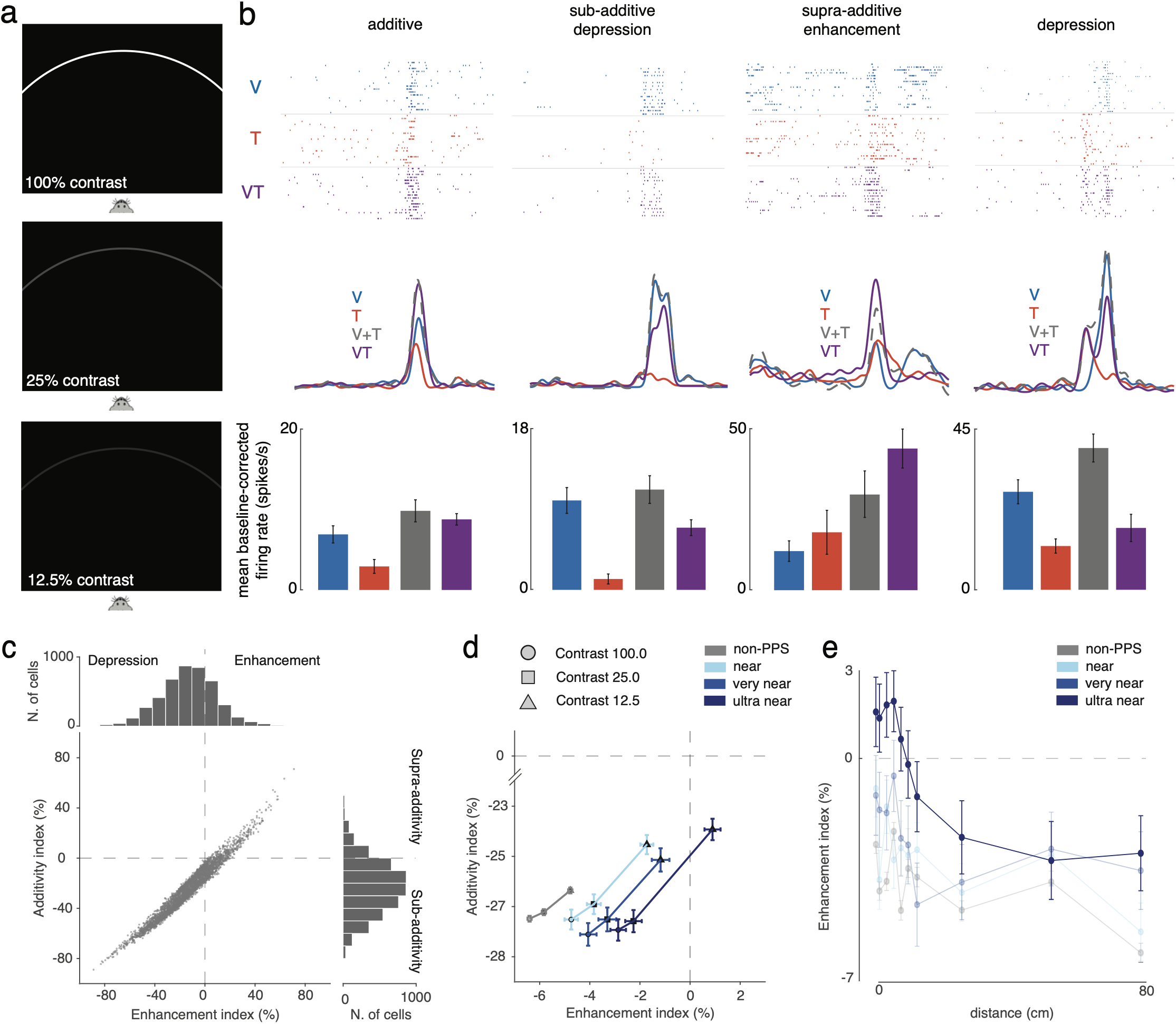
Peri-Personal Space Neurons Integrate Visuo-Tactile Signals. **A.** Schematic of the apparatus demonstrating the contrast manipulation (tactile and different visual distances not shown). **B.** Example neurons demonstrating additivity, sub-additivity, and supra-additivity. Top row are raster plots, second row are PSTHs, and third row is mean baseline-corrected firing rate (0-300ms post-stimulus onset). Error bars are SEM across trials. **C.** Scatter plot of enhancement index vs. additivity index (see **Methods**) for all VISrl neurons. **D.** Mean and SEM enhancement and additivity index as a function of contrast (shapes) and PPS field (determined in a separate routine). **E.** Mean enhancement index as a function of visuo-tactile distance (in the multisensory integration subroutine, x-axis) and PPS field categorization (PPS mapping subroutine). Error bars are SEM (data is across all contrasts).

Neurons showed varied responses during paired visuo-tactile presentations. Some (**Fig. 5B**, left-most column) were enhanced relative to the unisensory modality that most readily drove spiking activity (i.e., VT > max (V, T)) and best characterized as linear operators (or additive; VT = V + T). Others (**Fig. 5B**, second column) were depressed relative to the strongest unisensory responses, and demonstrated non-linear sub-additivity. Lastly, a relatively small fraction (see below) of neurons showed non-linear supra-additivity, with the paired response being greater than the sum of the unisensory constituents (**Fig. 5B**, third column). Lastly, while somewhat tangential to the current work, we also want to highlight that some neurons could be categorized across time first as linear, and then non-linear operators. For example, while visual and tactile stimulation occurred simultaneously, because of differential transmission times a paired response may follow first the tactile response linearly, and then the visual responses in a sub-additive and depressive manner (**Fig. 5B**, right-most column). Of course, these transmission/delay times will depend on the temporal relationship between external variables, but also visual spatial depth.

As a whole, the population of VISrl neurons is best categorized as showing multisensory depression (**Fig. 5C**, x-axis and marginal distribution, enhancement index < 0; see^55^) and sub-additive (**Fig. 5C**, y-axis and marginal distribution, additivity index < 0, see^55^). When splitting by contrast and whether the neuron indexed is a PPS neuron (and with either ultra-near, very near, or near spatial preferences based on the PPS mapping subroutine, see **Methods**) we observe evidence for both the “principle of inverse effectiveness” and the “spatial principle” of multisensory integration^64,65^. Namely, for all neuron types, lower contrast visual stimuli significantly increased both enhancement and additivity indices (all p < 0.001; **Fig. 5D**), in line with the principle of inverse effectiveness^64,65^. Across all contrasts, PPS neurons with ultra-near fields (< 5 cm) showed larger enhancement indices, followed by those with very near field (5-10 cm), then near fields (10-20 cm), and finally non-PPS neurons (ANOVA, p = 0.0006; follow-up corrected t-tests, all p < 0.05; **Fig. 5D**). This demonstrates the “spatial principle” of multisensory integration, in that by definition tactile receptive fields are on the body, and thus the overlap of tactile and visual receptive fields is largest for ultra-near PPS neurons, then very near, and finally near fields. Interestingly, field denomination (i.e., “ultra-near” vs. “very near”) seemingly had a greater impact on enhancement than additivity indices (interaction; p = 0.049). Only the “ultra-near” PPS fields during the lowest visual contrast presented showed on average (across visuo-tactile distances) a positive enhancement index (p = 0.024). No condition showed a positive additivity index on average.

We further examine this enhancement index as a function of visuo-tactile distance. This analyses demonstrated that PPS neurons with ultra-near fields showed multisensory enhancement up to 6cm from the body (all p < 0.05; **Fig. 5E**, across all contrasts). At 8 and 10cm, these neurons were neither enhanced nor depressed (both p > 0.38), and at 12.5cm and beyond, these neurons showed multisensory depression (p < 0.05). The other cell-types (i.e., very near and near PPS fields, as well as non-PPS neurons) always demonstrated multisensory depression (all p < 0.05).

Overall, the results demonstrate that PPS neurons do integrate visuo-tactile signals, following the spatial and inverse effectiveness “principles” of multisensory integration.

### Plasticity of Rostro-Lateral Visual Area Peri-Personal Space Neurons

We aimed to determine if PPS is a fixed map, or instead if and how VISrl PPS neurons would adapt to the statistical structure of their environment. Indeed, biologically-plausible neural network models and their corresponding experimental work^66^ suggest that PPS expands in humans after tool use because of repeated exposure to tactile stimulation on the body that occurs synchronously with visual stimulation occurring far from the body. Here, we therefore simulated a “tool-use” paradigm in rodents by exposing them to repeated touch during concurrent far visual cues. We first mapped PPS in a “pre” baseline condition, then exposed animals to repeated visuo-tactile stimulation in which touch was always delivered when a visual cue was presented far from the body, at 75 cm. We then mapped PPS again in a “post” condition. We recorded 4984 VISrl neurons across 24 sessions, 44 insertions, and 6 mice (see **Fig. S5B** for histological reconstructions).

We observed that PPS fields seemingly expanded in depth after repeated exposure to touch on the body during concurrent visual presentations far from the body. Indeed, firing rates of VISrl PPS neurons were reduced near the body (<15cm; p < 0.0005) and increased far from the body (>25cm; p < 0.0001) after visuo-tactile exposure (**Fig. 6A**). Examination of individual neurons suggested a degree of heterogeneity, with some PPS neurons not coding for visual depth after visuo-tactile exposure (**Fig. 6B**, top), some seemingly enlarging their receptive fields (**Fig. 6B**, middle), and some remapping, such that their near body preferences became a preference for farther distances (**Fig. 6B**, bottom). To quantitatively account for this heterogeneity, we performed k-means clustering, which determined there were two broad classes of neurons. First, 64.1% (SEM = 5.2%) did not remap as a function of visuo-tactile statistics (**Fig. 6C**, left). A second, smaller but still substantial group of neurons (35.9% ± 5.2%; χ² test, p = 0.007) seemingly demonstrated remapping, with peak visual responses drifting from ∼15-20cm “pre” exposure, to ∼35-40cm “post” exposure (**Fig. 6C**, right). Interestingly, it may be argued that the group remapping their receptive fields had a predisposition to do so, as their PPS fields in the “pre” condition already peaked farther than that of neurons that did not remap (p < 0.0001; **Fig. 6C**). Overall, this analysis suggests that PPS fields did not so much expand (**Fig. 6B**, middle example) than they did remap (**Fig. 6B**, bottom example) after repeated exposure to touch on the body and visual cues far from it. Further, the remapping was on the order of 20cm, or ∼20-25% the visuo-tactile distance repeatedly presented (75cm).

**Figure 6.**
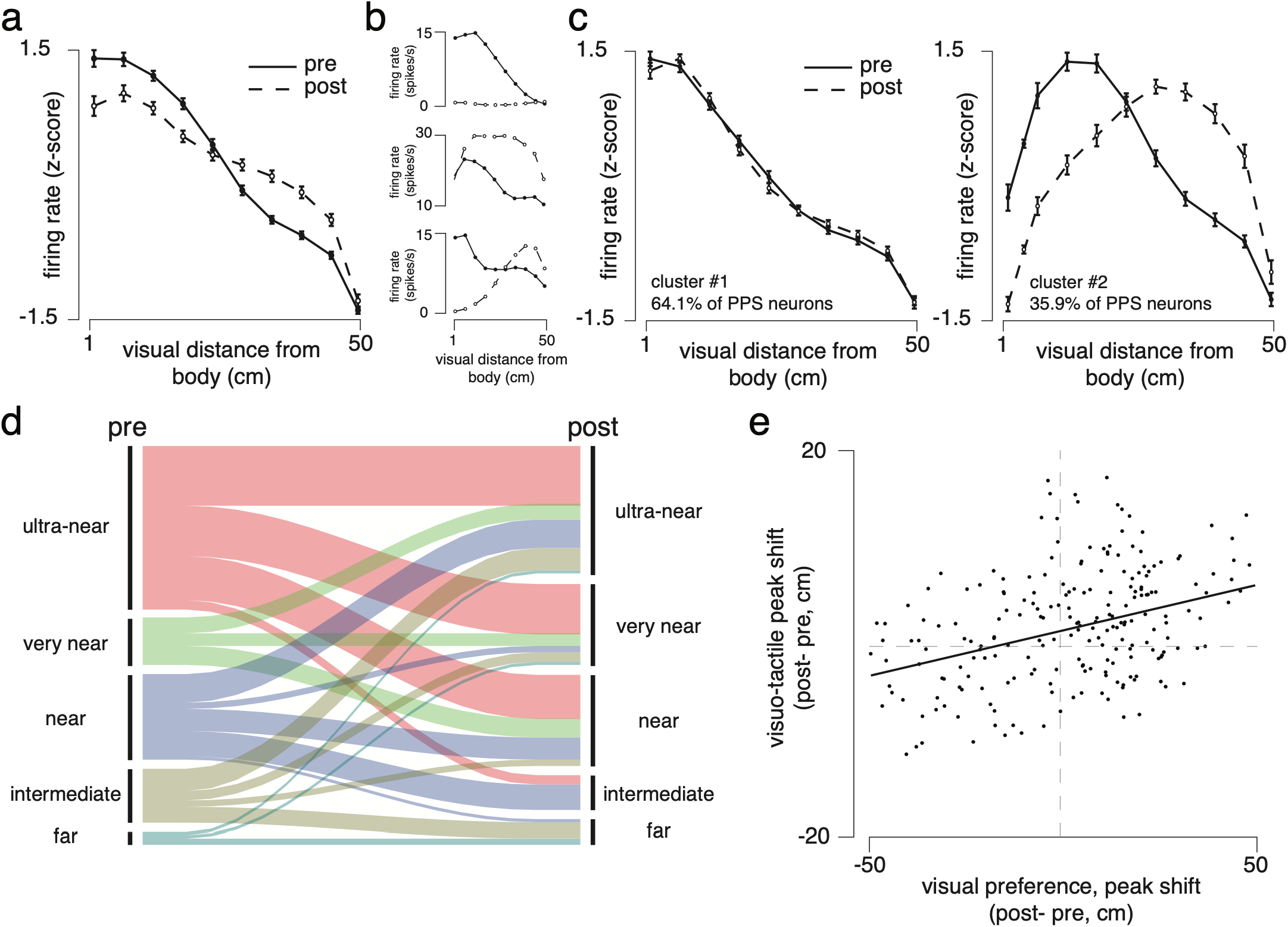
Plasticity of Rostro-Lateral Visual Area Peri-Personal Space Neurons. **A.** PPS fields pre (solid) and post (dashed) exposure to far visuo-tactile stimulation. Error bars are S.E.M. **B.** Three example visual tuning functions, respectively demonstrating the disappearance, enlarging, and remapping of PPS fields post exposure to far visuo-tactile stimulation. **C.** PPS fields pre (solid) and post (dashed) exposure to far visuo-tactile stimulation, separated by clusters that either did not (left) or did (right) remap. Error bars are S.E.M. **D.** Alluvial diagram showing the categorization of fields pre (left) and post (right) far visuo-tactile exposure. The length of black bars on either side show the frequency of each categorization. **E.** Correlation between changes in visual (post minus pre, x-axis) and visuo-tactile preference (post minus pre, y-axis) after exposure. These measures are correlated, yet field shifts are smaller for visuo-tactile facilitation than they are for visual tuning.

We categorized VISrl neurons as having ultra-near (< 5 cm), very near (5 – 10 cm), near (10 – 20 cm), intermediate (20 – 40 cm), or far (> 40 cm) fields, and examined how this categorization changed post far visuo-tactile exposure (**Fig. 6D**). Overall, these distributions shifted farther in space (see **Fig. 6D**, black bars “pre” and “post”; χ² test, p = 0.008) with the “ultra-near” denomination becoming less frequent after far visuo-tactile exposure, and the “very near” and “near” denominations becoming more common (χ² test, all p < 0.005). Indeed, while 37.9% (± 5.2%) of “ultra-near” fields remained post-exposure, 30.4% (± 5.1%) became “very near”, and 24.2% (± 6.7%) became “near” (**Fig. 6D**). The fraction of “far” fields also grew from pre- (3.5% ± 0.4%) to post-exposure (6.9% ± 1.1%, **Fig. 6D**).

Lastly, a similar effect was observed when categorizing PPS neurons not as responding to touch and having depth-dependent visual receptive fields (above), but as showing tactile response modulations as a function of visual proximity (**Fig. 3**). Overall, field remapping was significant but much smaller (2.33 ± 0.7 cm) when examining tactile facilitation (**Fig. S6**) rather than the preferred distance of visual tuning functions (17.4 ± 4.6 cm, p = 0.0036). The remapping of both metrics was correlated (r = 0.28, p < 0.001, **Fig. 6E**).

Overall, these results show that PPS fields remap as a function of visuo-tactile statistics, mimicking findings from macaques^25^ and humans^26,66^ during tool-use. We show that remapping occurs in about a third of neurons; cells that may already be somewhat pre-disposed to mapping in that their fields on average are already a step away from the body; and that this remapping is best characterized as a shift in receptive fields, and not an enlargement, though both occur.

### A “negative image” of grid cells?

In a final cohort of animals (Experiment 4; n = 6 mice, 20 sessions, 34 insertions, 6301 VISrl neurons; see **Fig. S5C** for histological reconstructions) we first identified PPS neurons (as above) and then examined their temporal frequency (TF), spatial frequency (SF), orientation, and contrast polarity preferences.

A subset of VISrl neurons were meaningfully driven by drifting gratings of different temporal frequency (3030/6301 or 48.09%, **Fig. 7A**), spatial frequency (13.3%), and orientation (16.5%; see **Methods**). The mean preferred temporal frequency of non-PPS VISrl neurons was 1.3 Hz (see^67,68^ for a similar estimate), with PPS neurons showing a preference for much faster moving gratings (8.4 Hz, p = 0.003; **Fig. 7B**, left). The mean preferred spatial frequency preference (at 10cm) was 0.032 cpd for non-PPS VISrl neurons, and slightly higher at 0.039 cpd (p = 0.041) for PPS neurons (**Fig. 7B**, right). Orientation preference was uniform for non-PPS VISrl neurons, while VISrl PPS neurons showed a preference for gratings implying motion toward their body (**Fig. 7C**, p = 0.002). Together, these results suggest PPS neurons are driven by coarse-grain, rapidly moving stimuli toward their body.

**Figure 7.**
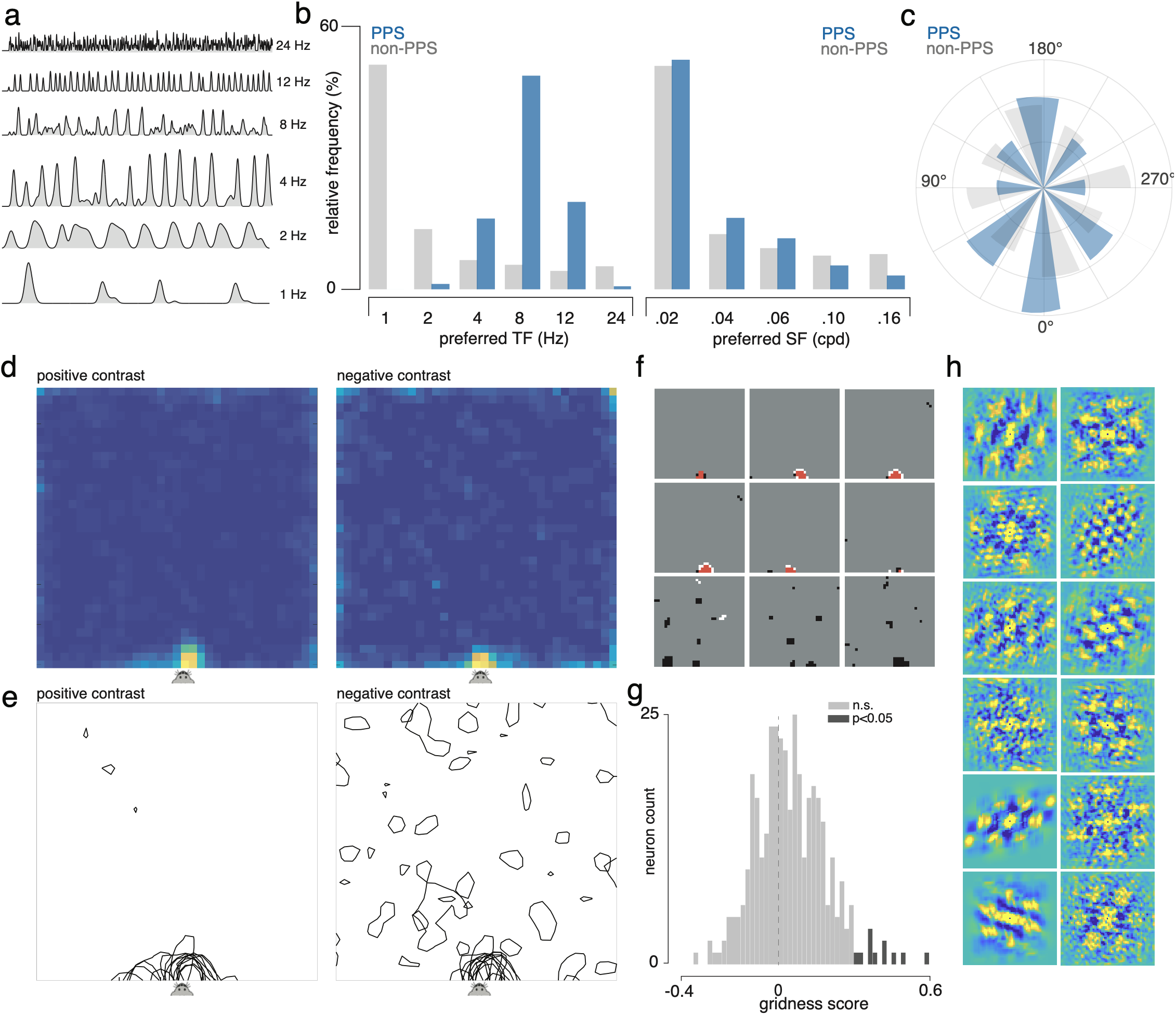
Temporal Frequency, Spatial Frequency, Orientation, and Contrast Selectivity. **A.** Example cell; firing rate as a function of time (4s) and temporal frequency of drifting grating. **B.** Preferred temporal (left) and spatial (right) frequency as a function of whether the cell was identifying in a separate routine as PPS or not. **C.** Preferred orientation with 0 degrees being toward the animal. **D.** Average firing rate map (70 x 70cm) as sparse noise white (left) or black (right) squares are presented. **E.** Example session, with firing rates now plotted as contour maps (z-score > 0.5). **F.** Example neurons, with their firing rate maps. Locations driven by white or black squares are depicted in their respective colors, while overlap between maps is shown in red. **G.** Distribution of gridness scores for the subset of neurons showing black fields at extra-personal distances. **H.** Example auto-correlations for neurons with a significant gridness score.

Sparse noise receptive field mapping suggested that on average PPS neurons were driven by both white (“positive contrast”; **Fig. 7D**, left) or black (“negative contrast”; **Fig. 7D**, right) cues within ∼4cm from the animal’s body, with no difference across these polarities. This PPS field size estimate is smaller than above, likely driven by the fact that here the visual stimuli have no motion (also see^69^). Interestingly, however, when we examined individual sessions (**Fig. 7E**), it appeared that in addition to very near fields (i.e., PPS), negative contrasts also frequently evoked responses at farther distances. Indeed, examination of individual PPS cells showed that while usually white and black contrast PPS fields overlapped (**Fig. 7F**, red), a subset of neurons were also driven by negative contrast cues at a multitude of locations. To examine if these “negative fields” had a spatial organization we computed their spatial autocorrelation and then a gridness score (see **Methods**)^70,71^. On average the population did not exhibit the hexagonal pattern characteristic of grid cell (0.004 ± 0.086, p = 0.11), but a small subset of neurons (29/317, 9.1% or almost twice the false positive rate expected given alpha = 0.05) did show a significant and positive gridness score (**Fig. 7G**, permutation testing). The auto-correlation of a subset of these neurons are shown in **Figure 7H**.

The results demonstrate that relative to non-PPS neurons, VISrl PPS neurons show a preference for faster, coarser grain stimuli moving toward their body. Further, they suggest that a minority of PPS cells may also demonstrate a “negative image” of grid cells in their extra-personal space.

## Discussion

We leveraged dense extra-cellular single neuron recordings to delineate a visuo-tactile egocentric map prioritizing peri-personal space (PPS) in the mouse rostro-lateral visual area (VISrl). This map extends beyond the reach of the whiskers: reaching tens of centimeters, whereas whiskers protrude only ∼1 to 3 cm past the snout at rest, and ∼3 to 5 cm during active whisking. Across the population of VISrl neurons, this map tiles the entire space from peri- to extra-personal when visual objects approach the body (up to 75 cm in our protocol). A recent study similarly identified VISrl as a site of distance-dependent visuo-tactile integration^69^. Using static object-location mapping, however, it described a representation largely confined to the ultra-near space the whiskers can reach, on the order of millimeters^69^. The far greater extent we report here likely reflects our use of dynamic stimuli, as our own estimates contract toward theirs when receptive fields are instead mapped with static sparse-noise probe.

This spatial PPS encoding is not a static representation. Instead, this map adapts dynamically as a function of the velocity of incoming stimuli, and plastically as a function of recent sensory statistics (e.g., paralleling a “tool-use” protocol). In this sense, it may be argued that VISrl PPS neurons encode a map of affordances^72,73^: the possible actions the environment “affords” agents (see^74^ for a similar argument in the hippocampus). Perhaps the most striking demonstration of this feature is that VISrl neurons seemingly track the entire trajectory of approaching cues (i.e., objects they may be able to interact with or which may be harmful), but not that of receding ones. Indeed, a receding object is one the animal can no longer act upon or be contacted by, and this brain area appears to stop representing these objects once they leave the actionable space. We argue, therefore, that PPS is not simply a passive, feedforward map, but likely an anticipatory representation of the space the body may soon interact with.

The current work extends beyond observations made in the seminal single-unit recordings that first delineated PPS in the macaque^4–13^. Namely, we show that PPS neurons are most often excitatory, and that they are functionally connected to their neighboring cells as a function of both physical proximity (following a Difference-of-Gaussians profile of near excitation, then inhibition, then excitation again) and tuning similarity. That is, PPS neurons show “like-to-like” connectivity^75,76^. The joint dependence on physical and feature proximity suggest that PPS coding follows a topographical map, a conjecture recently confirmed via imaging studies^69^. The near uniform inhibitory tone and like-to-like connectivity of excitatory neurons is also reminiscent of attractor networks, such as those seen in the head-direction system^77^. Moreover, PPS neurons prefer coarse spatial and fast temporal frequencies, consistent with VISrl PPS cells belonging to a cortical “where” pathway. The neurons that remap with sensory statistics appear predisposed to do so, in that at the outset they already encode distances one step removed from the body. As a population, the PPS network can remap to encompass both the animal’s physical body and the extra-personal space, yet individual neurons either do not remap with sensory statistics, or most often remap such that they no longer encode the body itself. Finally, we show that, at least in the mouse, PPS neurons integrate visual and tactile cues, most often sub-linearly, abiding by the spatial and inverse effectiveness principles of multisensory integration^64,65^. We further find that tactile responses of PPS neurons are facilitated by nearby visual cues, providing a neurophysiological foundation for the multisensory tactile facilitation long used to index PPS in humans^17,46–54^.

Visualizing time-resolved neural activity around the time of touch during concurrent approaching visual cues is particularly revealing as to the putative function and evolutionary origin of PPS neurons^3^. As one would predict for neurons that respond to touch and have visual receptive fields near the body, during visuo-tactile trials PPS responses are double-peaked: one response when touch is actually delivered, and a second when the visual object virtually “touches” the body. We conjecture that in evolutionary terms, virtual touch would most often have arisen from shadows, or from reflections on bodies of water. These experiences may have allowed early animals to learn appetitive and avoidant behaviors in the absence of real danger. Interestingly, in these evolutionary “virtual touches” the visual objects touching the body (e.g., shadows) would have been dark. This frames an unexpected finding. A small but meaningful subset of PPS neurons carried secondary receptive fields (**Fig. 7** and **Fig. 1D**, rightmost). These almost always responded to negative contrasts. Remarkably, some of these fields showed hexagonal tiling, reminiscent of grid cells^70–71^. That a neocortical visual area should carry the periodic, hexagonal signature long considered a hallmark of the hippocampal formation invites future work directed at comparing the canonical allocentric systems of the rodent brain with the egocentric, multisensory, plastic, and dynamic map uncovered here. Indeed, rodent spatial neuroscience has been read almost exclusively in allocentric terms, through grid cells^70, 71^, place cells^78^, head-direction cells^79^, and border cells^80^, among others. That machinery explains how an animal moves through the world. Yet it says little about how the world is represented relative to the body.

Egocentric codes are not absent near the hippocampus and throughout subcortical and midbrain networks. Object-vector cells signal the distance and bearing of objects^81^, and egocentric boundary cells appear in postrhinal and retrosplenial cortex^82,83^. Similarly, the superior colliculus (SC), a hallmark area for the study of multisensory integration^64,65^, sits at the center of a defensive circuitry putatively reading and interacting with PPS encoding. Indeed, separate collicular populations ultimately evoke flight or freeze: escape is driven largely by parvalbumin-positive SC neurons projecting to the parabigeminal nucleus, whereas freezing is mediated by SC neurons projecting to the lateral posterior thalamic nucleus^84,85^. Looming, on-collision stimuli tend to evoke flight, while distal, off-collision sweeps evoke freezing^86,87^. We speculate that VISrl is well positioned to inform this flight-or-freeze decision. By signaling whether and when an approaching object may make contact, it could bias behavior toward escape when impact is imminent, or toward freezing when a potential predator has not yet detected the animal. VISrl PPS neurons may therefore form a natural bridge between the hippocampal circuitry that supports navigation, and the subcortical and midbrain circuitry that drives defensive actions (see^88^). In the future, it will be interesting to more directly examine the role of PPS neuron in learning affordances, particularly those implying touch and derive from negative contrasts.

More broadly, the discovery of a visuo-tactile map prioritizing the rodent’s peri-personal space in neocortex opens a route to understanding circuit-level mechanisms guiding the interplay between the largely allocentric representations of the hippocampal formation and the egocentric, body-centered representations that guide perception and action.

## Acknowledgements

The authors thank Laurel Schuck and James Zhang for help with tissue clearing, imaging, and registration to the Allen CCF. This work was performed by the University of Minnesota Imaging Centers (RRID: SCR 020997) and the Minnesota Supercomputer Institute. This work was supported by the grant SFI-AN-NCSCN-00007276-10 from the Simons Foundation International as part of the SCENE collaboration, NIH R00NS128075, CTSI 1UM1TR004405, a Sloan Research Fellowship and a NARSAD Young Investigator Grant from the Brain & Behavior Research Foundation to JPN. The funders had no role in study design, data collection and analysis, preparation of the manuscript or decision to publish.

## Author Contributions

Conceptualization: JPN; Methodology: MJS, SB; Software: MB; Investigation: MJS, SB; Formal analysis: JPN, MB, EL; Visualization: JPN, EL; Resources: MB; Supervision: JPN; Funding acquisition: JPN; Writing – original draft: JPN; Writing – review and editing: all authors.

## Declaration of Interests

The authors declare no conflict of interest.

## Methods

### Animals

Experiments were performed in a total of 25 (n = 6 in Experiment 1; n = 7 in Experiment 2; n = 6 in Experiment 3; n = 6 in Experiment 4) male and female mice of mixed genetic background (C57BL6/j, JAX000664), between 11 and 33 weeks of age (17 ± 1.02 weeks). Animals were housed in a 12-hour reverse light/dark cycle. Male and female mice were first analyzed separately to assess potential sex-related differences. Given no differences were observed, male and female mice were grouped together for final analyses. All procedures performed in this study were approved by the Institutional Animal Care and Use Committee at the University of Minnesota (protocol 2405-42094A).

### Handling and Habituation

Mice were handled by experimenters for at least three days (increasing handling duration from 5 to 20 minutes) before any surgeries or experiments. After headbar implants but before craniotomies, they were also acclimated to head-fixation (3 days, increasing duration from 5 to 20 minutes).

### Surgeries

Each animal had two surgeries: a first one to secure a headbar on their skull allowing for head fixation, and a second one to perform craniotomies allowing for acute neurophysiology probe insertions.

For headbar implants, mice (∼10-14 weeks old) were initially anesthetized by placing them in an induction box at 3–5% isoflurane. They were then fixed in a stereotaxic frame via earbars and maintained at an anesthetized plane at 1–1.5% isoflurane. Under a microscope (Leica, M60), the dorsal surface of the skull was cleared of skin and periosteum, bregma and lambda were marked, the lateral and middle tendons were removed using fine forceps and the headbar was placed and cemented on a leveled skull. Cyanoacrylate (VetBond; World Precision Instruments) was applied to the edges of the skin wound to seal it off and minimize the risk of future infections. Finally, the exposed skull was covered with clear UV-curing optical glue (Norland Optical Adhesives 81; Norland Products). After surgery, mice were treated with carprofen for two days and given at least one week to recover before experiments.

For craniotomies, the induction procedures followed that of the headbar implants. Animals were affixed to stereotaxic frame via their headbars. The UV glue covering the skull was removed. On most animals, two craniotomies were made, targeting the left and right hemisphere VISrl at ± 3.25mm mediolateral and −2.75mm anteroposterior (negative ML values indicating the left hemisphere and negative AP values indicating posterior to Bregma). These craniotomies were performed with a dental drill (∼ 1mm in diameter) or a biopsy punch (1 mm in diameter). A small screw touching the brain was implanted for referencing. Craniotomies were covered with a low-viscosity silicon sealant (Kwik-Cast; World Precision Instruments) between days to prevent drying. Animals were given at least one day of recovery before starting neural recordings.

### Stimuli and Procedures – Experiment 1

The visual stimulus consisted of a full-contrast white annulus, 3 cm in width, presented on a black background. We used an annular stimulus because all points along the annulus are equidistant from the animal in depth, while this stimulus spanned a broad range of azimuthal visual angles. Across the range of depths tested, the stimulus subtended approximately from 46.05° to 180° of visual angle. On visual and visuo-tactile trials, the annulus moved either toward or away from the animal along the depth axis, at a speed of 50cm/s or 100cm/s. Visual stimuli were presented on a television monitor positioned nearly parallel to the ground (LG 55-inch OLED evo C4 Series; 122 × 70 cm; 1920 × 1080 pixels; 100 Hz refresh rate). The screen was placed at a 1.5° upward incline. Mice were head-fixed with the nostrils level with the vertical plane of the screen, and positioned at the midpoint of one of the short edges of the monitor. This geometry allowed visual stimuli to be presented up to 35 cm to the left and right of the animal, and up to 122 cm in front of the animal. The relative size of the mouse with respect to the screen is approximately equivalent to the relative size of an adult human on a basketball court. Tactile stimuli consisted of bilateral air puffs directed at the whisker pads from the front of the animal. Air puffs were 50 ms in duration and delivered at 1 bar, equivalent to 14.5 PSI. Compressed air was passed through a pressure regulator and routed to a three-way solenoid valve (P/N: LHDB1233418H; one inlet, two outlets). The two solenoid outputs were connected to 0.16 cm inner-diameter silicone tubes aimed at the left and right whisker pads. Tactile stimulus timing was controlled by a Bpod State Machine (r2.5), which sent TTL pulses to a Bpod Port Interface board to open the solenoid valve.

Stimulus presentation was controlled by custom-written Python code in PsychoPy (version 2025.1.0). Trial order was randomized across tactile-only, visual-only, and visuo-tactile conditions. Visual and visuo-tactile trials included approaching and receding annuli, different stimulus velocities (50cm/s and 100cm/s), and different visuo-tactile distances (1, 2, 4, 6, 8, 10, 12.5, 25, 50, or 75cm; i.e., distances of visual stimuli when touch was delivered). The experimental design included a full factorial of each of these conditions (e.g., visuo-tactile distances, visual speeds, and visual directions) and 60 repetitions per condition, for a total of 2700 trials per session. Inter-trial intervals were sampled randomly from a uniform distribution between 1.25 and 1.75 s.

### Stimuli and Procedures – Experiment 2

Experiment 2 was designed to characterize the multisensory integration properties of VISrl PPS neurons. The experiment consisted of two subroutines: a first to identify PPS neurons, and a second to characterize their multisensory integration properties. Importantly, both subroutines were conducted within a single neurophysiological recording session, allowing us to track individual neurons across both routines. The apparatus described for Experiment 1 was also used for Experiment 2. In the first subroutine, we identified PPS neurons using an experimental logic similar to that of Experiment 1. The only differences were that we tested a single visual stimulus direction (approaching) and a single velocity (125 cm/s). The visuo-tactile distances sampled were the same as in Experiment 1. Instead of 60 repetitions per condition, we performed 40 repetitions per condition, for each of a total of 21 conditions: 1 touch-only, 10 vision-only, and 10 visuo-tactile. Trial order was randomized, and intertrial intervals were randomly sampled from a uniform distribution between 1.10 and 1.40 s. These modifications were implemented to reduce the overall duration of the experiment. In the second subroutine, all sensory stimulation was brief and evoked, rather than dynamically moving. In touch-only trials, bilateral air puffs were delivered to the whisker pad at 1 bar for 50 ms. In vision-only trials, a 3 cm-wide annulus was presented for 50 ms at 1, 2, 4, 6, 8, 10, 12.5, 25, 50, or 75 cm from the mouse in depth. The visual stimulus was presented at full contrast, 100%, as in Experiment 1, or at 25% or 12.5% contrast. Visuo-tactile trials combined the touch-only and vision-only stimuli, with the two modalities always presented synchronously. Each condition was presented 18 times, in each of a total of 61 conditions: 1 touch-only, 30 vision-only, and 30 visuo-tactile. Trial order was randomized, and intertrial intervals were randomly sampled from a uniform distribution between 1.10 and 1.40 s. In total, each Experiment 2 session consisted of 1,938 trials.

### Stimuli and Procedures – Experiment 3

Experiment 3 was designed to test the malleability of VISrl PPS neurons by determining whether, and how, they remap after repeated sensory exposure indicating that touch occurs when visual stimuli are presented far from the body. This experiment consisted of three phases: a “pre” PPS mapping phase, an exposure phase, and a “post” PPS mapping phase. The pre and post phases were identical to the first subroutine used to map PPS fields in Experiment 2. Specifically, approaching visual cues were presented at 125 cm/s, and touch was delivered at 10 different distances from the body (max distance = 50cm). Touch-only and vision-only trials were also included. Each condition was repeated 40 times, across a total of 21 conditions. Trial order was randomized separately within the pre and post phases. During the intervening exposure phase, we repeatedly presented visuo-tactile trials in which touch was delivered when the visual stimulus was at the farthest distance from the body, 75 cm. This phase consisted of 1,520 identical trials. The same experimental setup was used here as in Experiments 1 and 2. Intertrial intervals were randomly sampled from a uniform distribution ranging from 1.10 to 1.40 s. In total, each Experiment 3 session consisted of 3,200 trials.

### Stimuli and Procedures – Experiment 4

Experiment 4 was designed to characterize the temporal frequency (TF), spatial frequency (SF), orientation, and contrast polarity preferences of VISrl PPS neurons. The experiment consisted of three subroutines, with continuous neural recording throughout. First, we identified PPS neurons using the first subroutine of Experiment 2. Next, we presented drifting gratings that varied in SF (0.02, 0.04, 0.06, 0.10, and 0.16 cycles/degree at 10 cm depth), TF (1, 2, 4, 8, 12, and 24 Hz), and orientation (0, 45, 90, 135, 180, 225, 270, and 315 degrees, with 0 degrees corresponding to straight ahead and motion toward the animal). Each condition (240 total) was presented at full contrast twice, for 4 s per presentation. Trial order was randomized. Finally, on a gray background screen, we presented either white or black patches, one at a time, within a 70 cm × 70 cm grid. Each patch was 2 cm × 2 cm and was presented 45 times for 20 ms, with trial order randomized. Intertrial intervals were randomly sampled from a uniform distribution ranging from 1.10 to 1.40 s.

### Neural Recordings

Neural recordings were performed with Neuropixels 1.0 probes^89^ (AP – 30kHz, gain = 500; LFP – 250Hz, gain = 250, IMEC), recording from the bottom third of electrodes (384 sites). Probes were mounted on a steel rod, which was held by micromanipulators (uMP-4, Sensapex). Probe trajectories were planned with Pinpoint^90^. Probes had a soldered connection to short the external reference to ground, and this latter one was connected to a screw fixed on the skull and in contact with the brain. The silicon sealant used to cover craniotomies was removed, and replaced with a silicone artificial dura repair compound (Dura-Gel; Cambridge NeuroTech). For later track localization, probes were labeled with CM-Dil (Thermo Fisher Scientific, V22888) by lowering the probe onto a coverslip or parafilm containing the dye (1 μl). The tips of the probes were maintained in CM-Dil until the dye dried out (∼20 s). On most recording days, we inserted two probes – one per craniotomy, at a 15° angle from vertical, and avoiding vasculature. The probes were lowered into position at ∼10 µm s^−1^. Electrodes were allowed to settle for ∼5-10 min before starting the recording. Data were acquired via a PXIe (PXI-1000; National Instruments) using SpikeGLX (Janelia Research Campus) and stored on a PC and cloud for subsequent analyses. At most, over four consecutive days we performed eight insertions in each animal (4 per craniotomy).

### Perfusions, Histology, and Probe Localization

Mice were deeply anesthetized with isoflurane and transcardially perfused through the left ventricle with phosphate-buffered saline, followed by 4% formaldehyde solution (Thermo Fisher Scientific, 28908). Brains were dissected, postfixed in the same fixative for at least 24 h at room temperature, washed, and stored in phosphate-buffered saline at 4°C for approximately 6–12 weeks. Whole-brain clearing was then performed using SmartBatch+ (LifeCanvas Technologies). Samples were washed in PBS on a 100 rpm shaker at 4°C for at least 4 days, with PBS replaced twice daily. Samples were then SHIELD-fixed, incubated in 20 mL Delipidation Buffer on a 100 rpm shaker at 45°C for 24 h, transferred to mesh bags, and actively cleared in the SmartBatch+ device at 42°C, 40 V, and 1250 mA for 30 h in 40 mL Delipidation Buffer. Cleared samples were incubated in 20 mL of 50% EasyIndex in deionized water on a 100 rpm shaker at room temperature for 24 h, followed by 20 mL of 100% EasyIndex (RI = 1.52) for at least 3 days before imaging. Whole brains were imaged using an AxL Cleared Tissue LightSheet microscope (3i) at 4 μm isotropic voxel resolution. Imaging was performed in EasyIndex (RI = 1.52) at 30°C using 5×/0.14 NA excitation objectives and a 1.0×/0.25 NA collection objective with 1.63× zoom. Autofluorescence was imaged with 488 nm excitation and a 525/50 emission filter, and DiI-labeled electrode tracts were imaged with 561 nm excitation and a 630/92 emission filter. Both channels were acquired with a 200 ms exposure. Next, images were downsampled to 25 × 25 × 25 μm voxel resolution to match the Allen Mouse Brain Atlas (CCF), rotated into the target LSP orientation, and registered to the Allen Atlas using ANTsPy. Registration was performed by applying affine followed by nonlinear deformable transformations, optimizing mutual information between the sample images and the atlas. Lastly, we reconstructed the location of probes by manually tracing the fluorescent dye on CCF-aligned coronal and sagittal images using a Python-based image viewer (Lasagna) equipped with a plugin tailored for this task. Finally, we manually aligned electrophysiological and anatomical landmarks along the probe trajectory using a custom histology-to-electrophysiology alignment tool.

### Spike Sorting, Curation, and Quality Control

Sensory stimuli (e.g., location of annulus) and neural data were synchronized by logging and sending to each acquisition device a pseudo-random barcode, TTLs of random duration between 1 and 5 seconds. These traces are then cross-correlated across devices, with their intercept expressing the delay between recording devices and the slope of this regression (always >0.99) expressing the clock drift. These are then used to synchronize spiking activity to the timing of sensory stimuli presentation.

Neural data were spike sorted with Kilosort 4^91^, using default parameters and the appropriate channel map. After automatic sorting units underwent three steps of quality control: a first manual step, a second semi-automated step and a final automated one. First, we manually inspected clusters in Phy for noise artifacts. Units with non-biological waveforms were discarded. Then, we applied the Recording Inclusion Guidelines for Optimizing Reproducibility (RIGOR) criteria developed by the International Brain Lab^92^. These include (1) inspecting entire sessions by constructing ‘drift maps’ (channel × time, spikes as dots) and establishing no severe tissue or electrode drift, (2) a yield of at least 0.1 single neurons per electrode channel in each brain region, (3) a median action potential band RMS less than 40μV, (4) a median derivative of 20–80 Hz LFP-band power less than 0.05 dB/µm, and (5) the absence of epileptiform activity. If these metrics were not met, the entire session was discarded. At the level of single units, neurons had to satisfactorily pass (6) a sliding refractory-period metric estimating less than 10% contamination, with greater than 90% confidence, across possible refractory periods from 0.5 to 10ms, (7) and have median spike amplitude greater than 50 µV, among others (see^93^ for more detail). Lastly, we performed automated classification of units as “good” single units by applying the default criteria from Bombcell^94^.

### Analyses – Experiment 1

#### Single unit analyses

A unit was classified as tactile-responsive if its firing rate differed between a baseline window (−500 to −50 ms) and a response window (+50 to +500 ms) relative to air-puff onset (Wilcoxon signed-rank test, p < 0.05). All subsequent analyses were restricted to tactile-responsive VISrl units. Then, for each neuron we computed firing rate as a function of visuo-tactile distance, separately for each motion direction and speed. Tuning to distance was quantified as the mutual information between firing rate and distance,

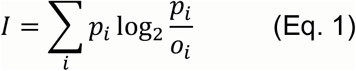

where *p_i_* is the trial-averaged firing rate at distance bin *i* normalized across the N bins to a probability distribution ∑*_i_ p_i_* = 1, and 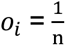 is the (uniform) occupancy. With uniform occupancy this is equivalent to the Kullback–Leibler divergence of the tuning curve from a uniform distribution. Significance was determined by a permutation test in which distance-bin labels were shuffled within trials (1000 shuffles). A neuron was deemed distance-tuned when its observed information exceeded the 95th percentile of the null distribution (p < 0.05). To quantify how stimulus speed reshaped distance tuning, we fit each neuron’s firing rate as a function of distance with a four-parameter sigmoid (fit by nonlinear least squares),

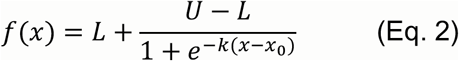

where L is the lower asymptote, U is the upper asymptote, *x*_0_ is the midpoint or inflection point, and k controls the steepness of the curve. The inflection point *x*_0_ indexed the distance separating near from far space, and the slope k indexed the sharpness of the near-to-far transition. We compared *x*_0_ and k across motion directions (approach, recede) and speeds (50, 100 cm/s). Shifts in the distribution of preferred distances across speed conditions were tested with chi-square tests. For this analysis, we only retained neurons with a sigmoidal fit with r^2^ > 0.75. Tactile-responsive, distance-tuned neurons were initially classified as classically-defined PPS neurons when their visual distance-tuning function peaked within 20 cm of the body (near preference; **Fig. 1**). This near-preference criterion was used only for the classical definition and was relaxed in data-driven clustering (**Fig. 2**). Namely, neurons were clustered according to the shape of their visual distance-dependent response profiles using k-means clustering after z-scoring each neuron’s tuning curve across distances. The number of clusters was selected automatically by fitting models across a range of candidate k values (2 to 10) and choosing the solution with the highest mean silhouette score, which quantifies how well each neuron fits within its assigned cluster relative to other clusters^95^. For visualization only, the z-scored tuning functions were projected to two dimensions with t-SNE^59^. Lastly, for neurons with tactile responses modulated by visual distance, we computed, per visuo-tactile distance, the evoked response as the spike count in the response window (+50 to +500 ms after tactile onset) minus the baseline window (−500 to −50 ms), and subtracted the matched tactile-only evoked response. Facilitation as a function of distance was tested with a one-way ANOVA across distances, with post hoc paired t-tests of each distance against zero (modulation relative to tactile-only).

#### Waveform-based classification of putative excitatory and inhibitory neurons

For each well-isolated unit we extracted the mean spike waveform on its peak channel and computed the trough width of the extracellular action potential. We classified units with narrow waveforms (trough width < 0.35 ms) as putative inhibitory neurons and units with broad waveforms (trough width > 0.45 ms) as putative excitatory neurons, following the waveform-based separation of fast-spiking inhibitory from regular-spiking excitatory cells used in sensory cortex (see^60^). Units with intermediate trough widths were left unclassified. Fractions of putative excitatory and inhibitory neurons were computed per session and are reported as the mean ± S.E.M. across sessions.

#### Inference of putative monosynaptic connectivity

To infer putative monosynaptic interactions between simultaneously recorded VISrl-PPS neurons, we applied a generalized linear model for cross-correlation analysis (GLMCC^61^). For each ordered pair of neurons, we constructed the cross-correlation function of their spike times over lags of −50 to +50 ms at 1 ms resolution. The GLMCC fit isolates short-latency, monosynaptic coupling from slower co-fluctuations in firing rate by modeling the cross-correlogram as the sum of a slowly varying component and a fast post-synaptic coupling term, the sign of which defines the interaction as excitatory or inhibitory. A putative connection was retained only when (1) the inferred coupling exceeded the confidence interval for the null hypothesis of no connection, and (2) a pre-trained convolutional neural network independently classified the pair as a putative monosynaptic connection^62^. We summarize the fraction of retained connections classified as excitatory, and the agreement between waveform-based and spike-timing-based cell-type classifications, across sessions. We then asked how putative connectivity depended on the separation between neurons in physical and feature space. For each session we assembled a signed connectivity matrix (excitatory > 0, inhibitory < 0) and computed connection probability as the number of connected pairs divided by the number of possible pairs within a given separation (self-pairs excluded). Physical separation was the pairwise Euclidean distance between estimated soma positions. Feature-space separation was the absolute difference in cluster assignment, where clusters were defined by visual spatial tuning and ordered by preferred visuo-tactile distance.

#### Population dynamics

We computed state-space trajectories using demixed principal component analysis (dPCA^63^). Demixed PCA decomposes population activity into components that, unlike standard PCA, remain interpretable with respect to task variables. We marginalized neural activity over four factors, visual velocity, visual direction, visuo-tactile distance, and time, together with their interactions terms. All epochs were aligned to tactile stimulus onset (t = 0). The condition-independent (time) marginalization captures the tactile-evoked response common across conditions. We retained the first 20 components and report, for each marginalization, the fraction of total population variance it explained, alongside the variance captured by standard PCA over the same number of components.

### Analyses – Experiment 2

#### PPS neuron definition

Using the separate PPS mapping subroutine in the same session, a neuron was classified as a PPS neuron if it was tactile-responsive (two-sided Wilcoxon signed-rank test against zero, p < 0.05) and significantly modulated by visual distance (Kruskal-Wallis test of the visual response across the ten sampled distances, p < 0.05). Each PPS neuron’s preferred distance was taken as the distance of its peak mean visual response and labeled by proximity, ultra-near (< 5 cm), very-near (5–10 cm), or near (10–20 cm). Non-PPS neurons served as the comparison group.

#### Response quantification

For every trial we computed a baseline-corrected evoked firing rate, defined as the mean rate in a post-stimulus response window minus the mean rate in a pre-stimulus baseline window (baseline −0.50 to −0.10 s relative to stimulus onset). Two response measures were computed and gave consistent results: a fixed-window spike-count rate (response window 0.10 to 0.50 s) and a peak-PSTH rate (mean rate in a ±50 ms window centered on each neuron’s response peak, where the peak time was the maximum of the smoothed PSTH built from all pooled multisensory trial events, 10 ms bins, Gaussian-smoothed). Each condition required at least forty trials. In the main text, we use the spike count metric.

#### Integration indices

For each neuron, and separately for each contrast and visuo-tactile distance, we quantified multisensory integration following the additive and enhancement frameworks of Avillac et al. (2007). Let *C*, *V*, and *T* be the baseline-corrected evoked responses on combined visuo-tactile, visual-only, and tactile-only trials. We computed an enhancement index, comparing the multisensory response to the stronger unisensory response,

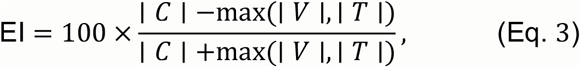

and an additivity index, comparing the multisensory response to the linear sum of the unisensory responses,

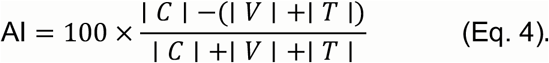

Positive EI indicates multisensory enhancement relative to the best unisensory input, positive AI indicates supra-additivity, negative AI indicates sub-additivity. Note, a neuron can be sub-additive but still demonstrate enhancement. Each neuron’s integration type (EI, AI) was classified by a nonparametric bootstrap (500 resamples of trials, with replacement, within each modality). On each resample we computed the deviation from the linear sum, C - (V+T), and the deviation from the best unisensory response, C - max(V,T), yielding two-sided bootstrap p-values. Neurons were classified as supra-additive (C > V+T, p < 0.05), sub-additive (C < V+T, p < 0.05), enhanced (C > max(V,T), p < 0.05), or depressed (C < max(V,T), p < 0.05), and as additive otherwise.

### Analyses – Experiment 3

Analyses were restricted to VISrl PPS neurons, defined as units that were tactile-responsive and significantly tuned for visual distance in the pre-exposure (“pre”) condition. A neuron was classified as visually distance-tuned when a one-way ANOVA of firing rate across distance bins reached p < 0.05. Further, to assure these neurons were stable, we also filtered neurons to have an effect size of η^2^≥0.02. Parallel criteria (ANOVA p < 0.05, η^2^≥0.02) were applied to defined neurons showing a tactile modulation during visuo-tactile stimulation.

#### Visual distance tuning

Visual responses were expressed as a continuous function of stimulus depth. For each pair of consecutive stimulus frames within a trial we computed an instantaneous firing rate,

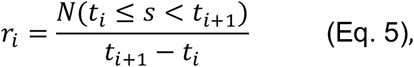

where N(⋅) is the spike count in the inter-frame interval. Firing rates were binned into 5 cm distance bins, and bins with fewer than three hundred samples were excluded. Tuning was tested with a one-way ANOVA across bins, with effect size

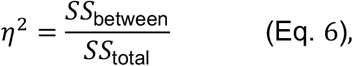

and the preferred distance was taken as the bin with the maximum mean firing rate. Each neuron’s preferred distance was labeled categorically as ultra-near (< 5 cm), very-near (5–10 cm), near (10–20 cm), intermediate (20–40 cm), or far (≥ 40 cm).

#### Pre versus post comparison

The identical procedure was applied to the post-exposure (“post”) recordings. For each neuron we built matched pre and post tuning curves on a common distance axis and quantified remapping as the change in preferred distance,

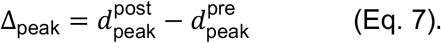

Transitions between pre and post distance categories were tabulated and tested with χ² tests on the category-count distributions.

#### Clustering of remapping profiles

To classify the heterogeneity of post-exposure tuning shapes without imposing a prior, each neuron’s post tuning curve was z-scored across distance and submitted to k-means clustering (squared-Euclidean distance, up to 1000 iterations). The number of clusters was chosen objectively by maximizing the mean silhouette value over k = 2 to 10; if no solution exceeded a mean silhouette of 0.10 the population was treated as a single cluster. This procedure identified two reliable classes, interpreted as a non-remapping (**Fig. 6C**, left) and a remapping population (**Fig. 6C**, right). The pre-exposure preferred distances of the two classes were compared to test whether remapping neurons were predisposed by an already more distal baseline tuning.

#### Combined visuo-tactile facilitation

In parallel, in combined visuo-tactile trials we characterized each neuron’s tactile response as a function of the concurrent visual distance. For each tactile stimulation delivery we computed a baseline-subtracted evoked firing rate from the trial PSTH (baseline −0.50 to −0.10 s, response window 0.10 to 0.50 s around touch onset, 10 ms bins, Gaussian-smoothed), averaged across trials at each sampled position, and tested for spatial modulation with a one-way ANOVA (preferred distance = peak position). Pre-to-post remapping of this metric was computed as for the visual tuning, and the two forms of remapping (visual preferred-distance shift and combined-modulation shift) were related across neurons by Pearson correlation.

#### Reliability

To confirm that pre/post shifts were not driven by within-session instability, tuning curves were recomputed on split halves of the trials (first vs second half, or odd vs even) within each phase and condition, and the resulting split-half preferred-distance estimates were compared to the across-phase shifts.

### Analyses – Experiment 4

#### Drifting-grating tuning analysis

For each neuron we characterized visual responses along three stimulus dimensions: temporal frequency (TF), spatial frequency (SF), and orientation. Responses were quantified from the peri-stimulus time histogram *r(t)* of each 4 s grating presentation using the first two Fourier components, following the framework of Skottun et al. (1991)^96^. The mean evoked rate (the zeroth harmonic, F0) captures the unmodulated response (or elevation of response),

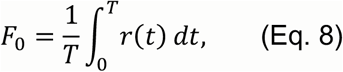

while the first harmonic (F1) captures the component of the response modulated at the grating’s drift frequency (i.e., modulation depth of the response), *f*_TF_

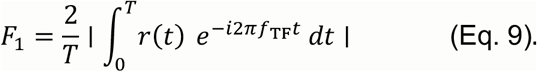

F0 reflects how much a neuron fires on average, whereas F1 reflects how strongly its firing is entrained to the passage of successive light and dark bars, so a neuron whose spike trains track the grating cycle produces a large F1 even when its mean rate is modest. For each dimension *d* ∈ {TF, SF, orientation} we tested for tuning with a one-way ANOVA on the F1 response across the levels of *d*, holding the other two dimensions at the neuron’s preferred value, and defined the preferred value as

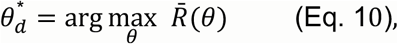

where *R̄* (*θ*), is the mean response at level *θ*. A neuron was called tuned for dimension *d* when its ANOVA reached p < 0.05.

#### Sparse-noise receptive-field analysis

VISrl neurons were probed with a sparse-noise stimulus in which a single white (+1) or black (−1) square (2cm x 2cm) appeared at a pseudorandom location on a 35 × 35 grid (70 cm wide x 70 cm depth), refreshing every 20 ms. For every grid location and contrast, we computed a trial-wise change in firing rate between a post-stimulus window (50–300 ms after square onset) and a pre-stimulus baseline (−300 to −100 ms),

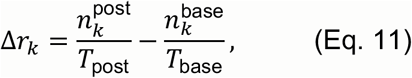

where *n_k_* is the spike count on presentation k and T the window duration. Per pixel we summarized the response with a one-sample t-like statistic across presentations,

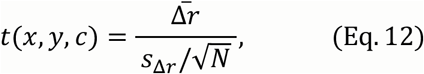

yielding separate 35 × 35 maps for white and black squares. Maps were lightly smoothed (Gaussian, σ = 1 pixel), thresholded at ∣t∣ ≥ 2.0.

#### Gridness score analysis

To test whether a neuron’s visual response field exhibited the hexagonal periodicity characteristic of grid cells^70^, we computed a gridness score from its 2D sparse-noise response map (analyzed only for black squares, as white did not produce extra-personal fields). Following the spatial-autocorrelation approach of Sargolini et al. (2006)^71^, we first formed the map’s full 2D autocorrelogram as,

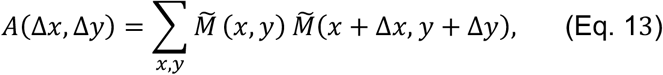

where 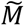 is the response map after removing its mean (DC offset), and normalized *A* by its peak absolute value. We then isolated an annular region of the autocorrelogram centered on the central peak (inner radius 3 px, outer radius set to just inside the map edge), excluding the central peak itself. The annulus was rotated to 30°, 60°, 90°, 120°, and 150° and each rotated version was correlated (Pearson) with the unrotated annulus, giving rotational correlations *C*_θ_. The grid score was defined as the difference between the correlations at the grid-consistent angles and the grid-inconsistent angles,

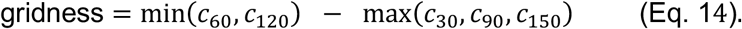

A hexagonal lattice produces autocorrelation peaks every 60°, so it ought to correlates highly with itself at 60° and 120° but poorly at 30°, 90°, and 150°, yielding a positive score (theoretical range roughly −2 to +2). Maps with fewer than ten finite pixels, flat maps, or annuli too small to evaluate were assigned an undefined score and excluded. This procedure was also applied after circular shifting of spikes for permutation testing (1000 shuffles).

## Supplementary Materials

**Figure S1.**
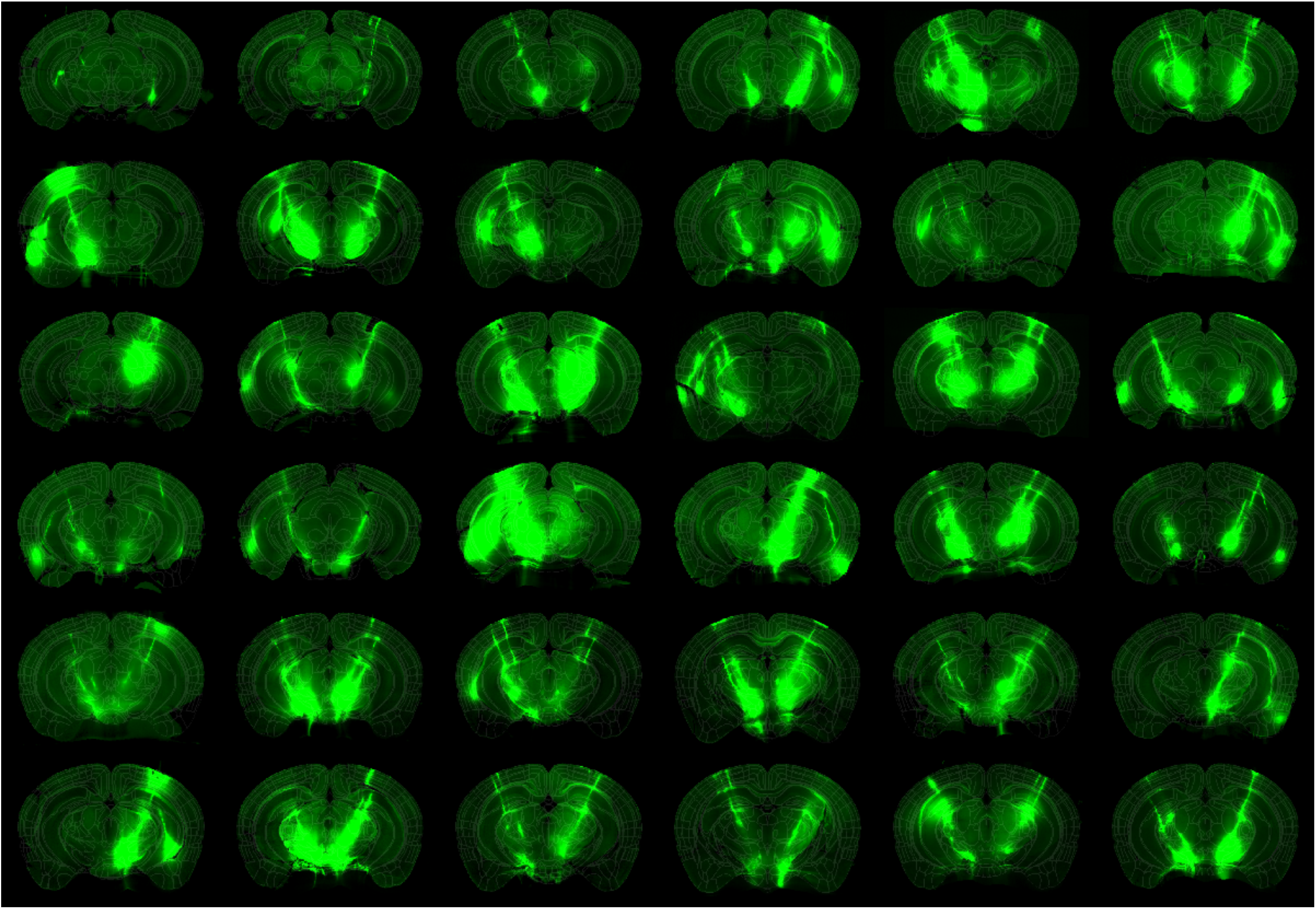
Histological reconstruction of probe locations. Example brain slices (n = 36) where we have visualized the dye (green) and the Allen CCF atlas is super-imposed in white lattice.

**Figure S2.**
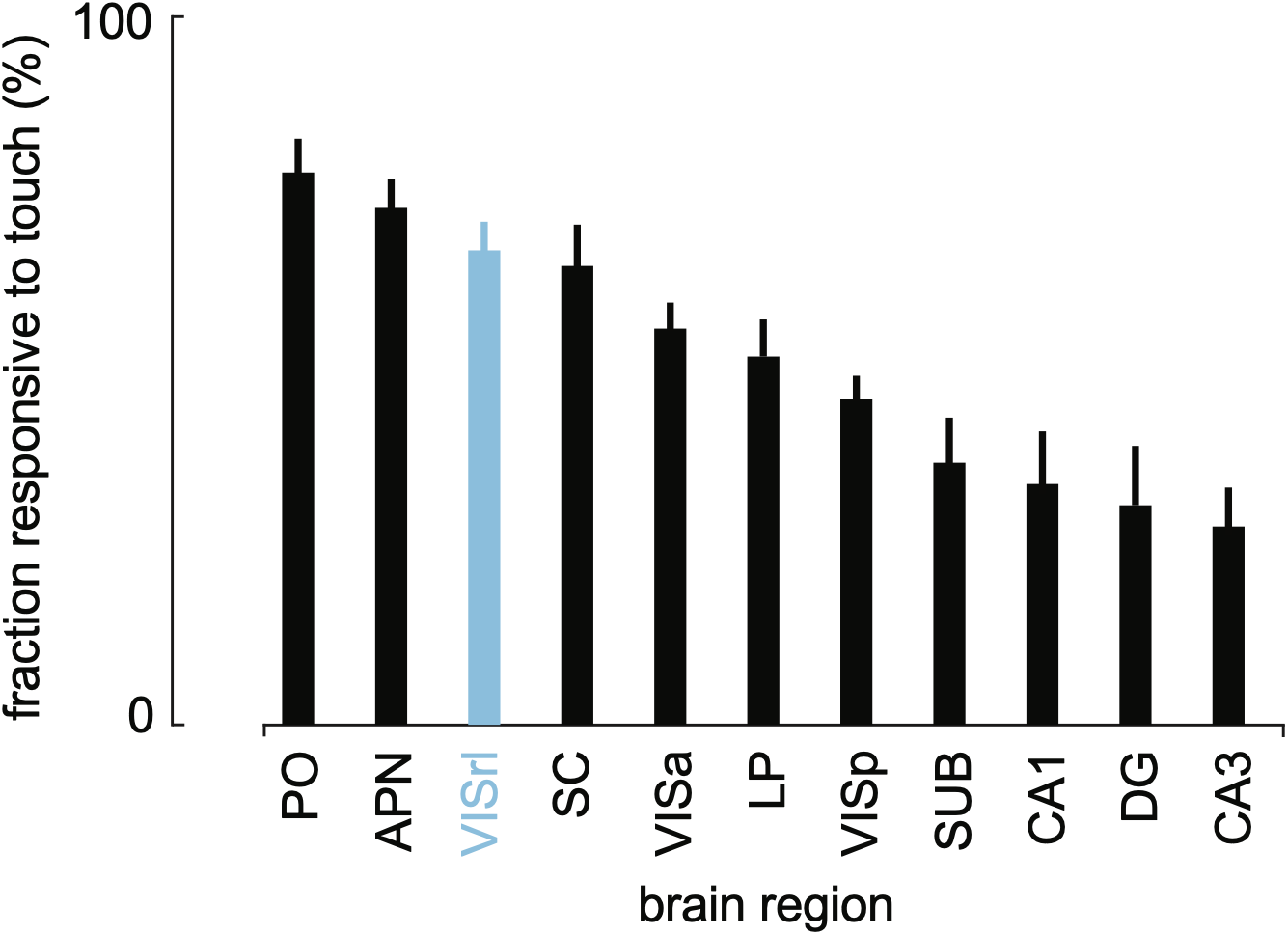
Fraction of neurons responding to tactile onset as a function of brain area. In order; the secondary whisker somatosensory thalamus (PO), the anterior pretectal nucleus (APN), rostro-lateral visual area (VISrl), the superior colliculus (SC), anterior area of the mouse visual cortex (VISa), the lateral posterior nucleus of the thalamus (LP), primary visual cortex (VISp), Subiculum (SUB), field CA1, dentate gyrus (DG), and field CA3. Bars are means across sessions (minimum number of neurons = 200, minimum number of sessions 4), and error bars are S.E.M.

**Figure S3.**
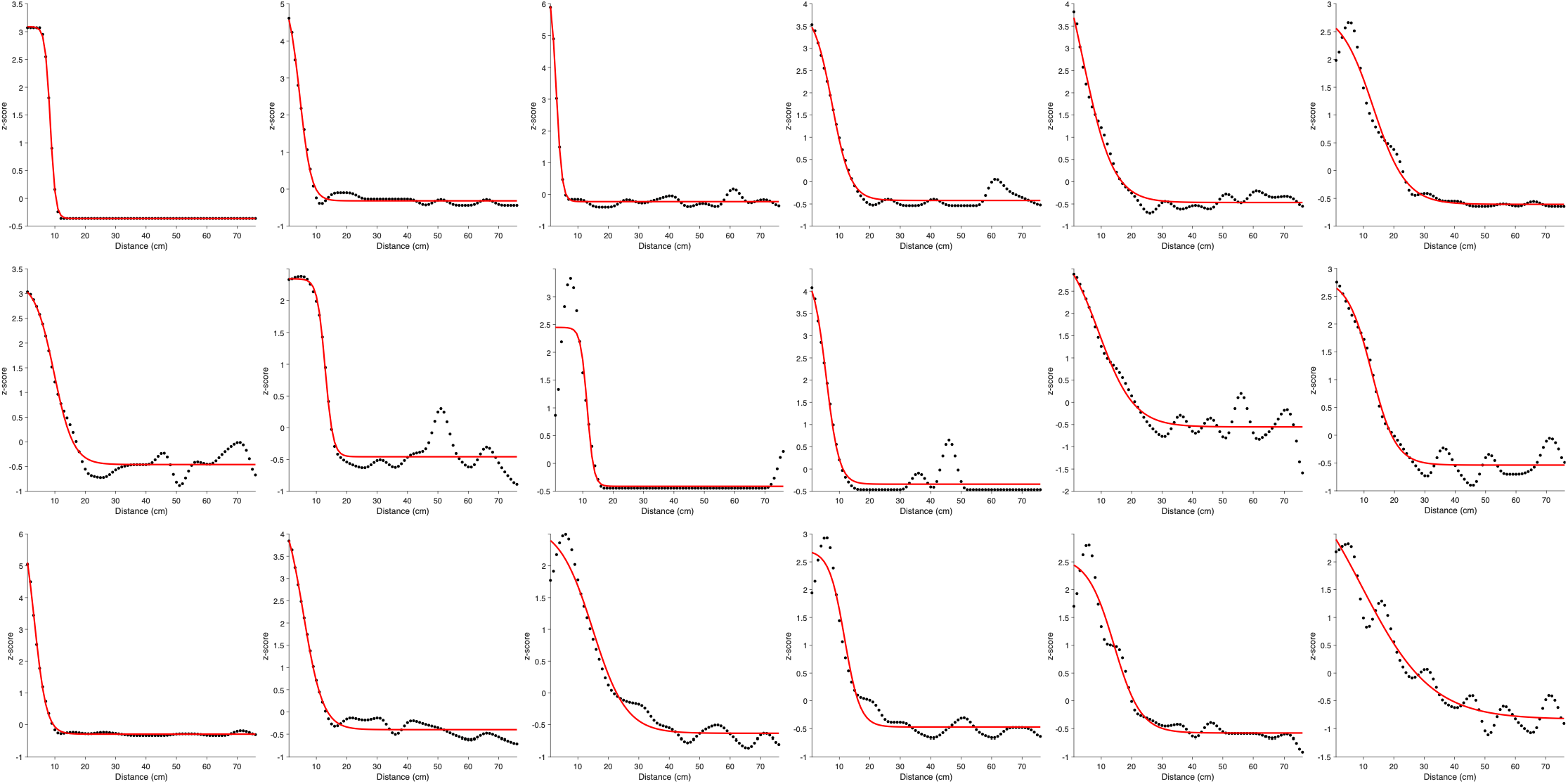
Example visual tuning function fit to sigmoidal curves. Firing rates as a function of distance (black dots) are fit to a sigmodal curve (red) in order to extract their central point and slope. Examples here were chosen at random, and include examples from all directions and velocities.

**Figure S4.**
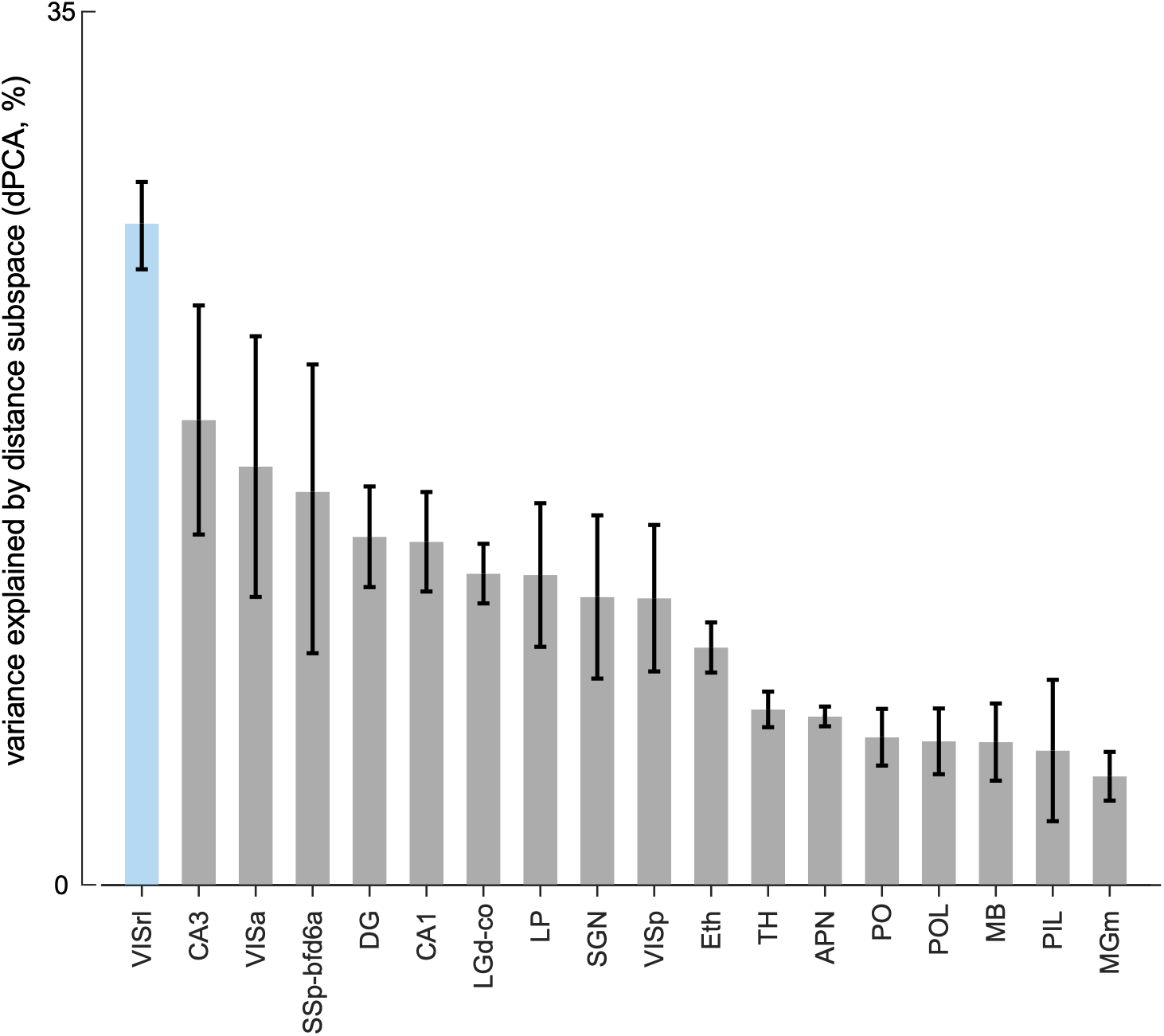
Variance explained by the visuo-tactile ‘distance’ subspace in different brain areas. Brain areas are included only if they have at least.2 sessions with at least 20 neurons recorded from simultaneously. Error bars are Standard Error of the Mean (SEM).

**Figure S5.**
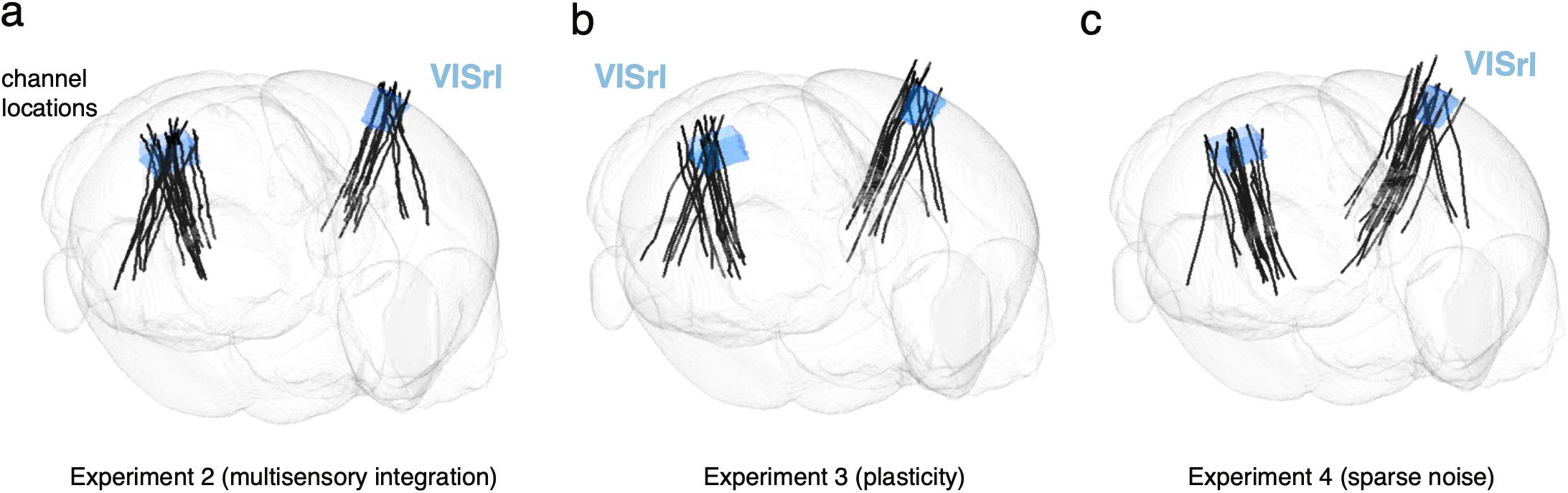
Probe location reconstruction for Experiments 2-4.

**Figure S6.**
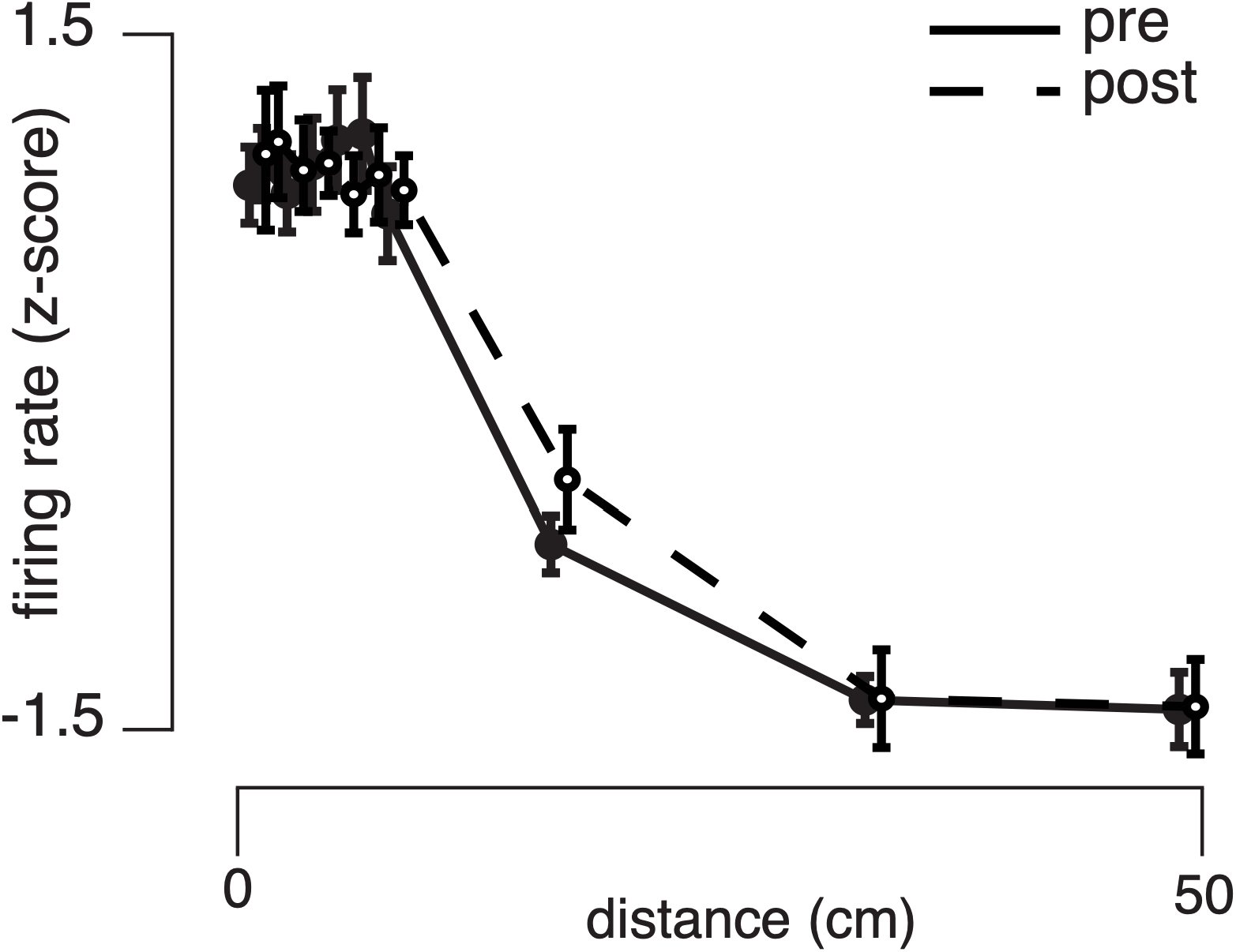
Remapping of PPS neurons defined as modulation of tactile response as a function of visuo-tactile distance. PPS fields pre (solid) and post (dashed) exposure to far visuo-tactile stimulation. Error bars are S.E.M.

## Notes

### Competing Interest Statement

The authors have declared no competing interest.

